# Structural basis of a high-affinity antibody binding to glycoprotein region with consecutive glycosylation sites

**DOI:** 10.1101/2022.07.24.501275

**Authors:** Yunbin Han, Jun Niu, Deng Pan, Chunchao Feng, Ke Song, Bing Meng, Ulrika Westerlind, Yan Zhang, Haiguang Liu, Lan Xu, Dapeng Zhou

## Abstract

Consecutive glycosylation sites occur in both self and viral proteins. Glycan-shielding of underneath peptide region is a double-edged sword, that avoids immune attack to self-proteins, but helps viruses including HIV-1 and SARS-CoV2 to escape antibody binding. Here we report a high-affinity antibody, 16A, binding to linear peptide containing consecutive glycosylation sites. Co-crystallization of 16A Fab and glycopeptides with GalNAc modifications at different sites showed that STAPPAHG is the sequence recognized by 16A antibody. GalNAc modification at Threonine site on STAPPAHG sequence significantly increased the affinity of Fab binding by 30.6 fold (KD=6.7nM). The increased affinity is conferred by hydrophilic and pi-stacking interactions between the GalNAc residue on Threonine site and a Trp residue from the CDR1 region of the heavy chain. Furthermore, molecular modeling suggested that GalNAc on T site causes more favorable conformation for antibody binding. These results showed that glycan modification most proximal to linear peptide core epitope significantly increases antigenicity of a glycopeptide epitope. The antibody recognition mode by peptide-binding CDR groove with a glycan-binding edge, may shed light on designing of linear glycopeptide-based vaccines for cancer and viral diseases.

**Teaser:** A high-affinity antibody was found to bind densely glycosylated glycoprotein region by a peptide binding groove of the antibody’s variant region, with a glycan-binding edge specific to glycosylation site most proximal to core peptide epitope.

## Introduction

Dense glycosylation is found in glycoproteins with critical biological functions.Mucins are a family of heavily-glycosylated proteins with critical roles in the transcriptional regulation of genes associated with tumor invasion, metastasis, angiogenesis, proliferation, apoptosis, drug resistance, inflammation, and immune regulation (*1*). Viruses hijack the glycosylation of host cells to avoid immune attack, as exemplified by HIV Env protein gp120 (*2, 3*) and Spike protein of SARS-CoV2 (*4, 5*). Vaccine strategies targeting densely glycosylated peptides have been studied in past decades (*6, 7*). Antibodies were reported to target the glycan portion alone, the peptide alone, or both glycan and peptide portion (*8*). Based on epitope mapping of antibodies, linear Mucin 1 peptide SAPDT(αGalNAc)RPAP was first tested as cancer vaccine (*6*), and reported as superior in tumor prevention as compared to non-glycosylated peptide vaccines.

Mucin 1 (MUC1) has been ranked No. 2 of all 75 tumor-associated antigens as cancer vaccine targets evaluated by National Cancer Institute Translational Research Working Group, based on certain criteria, such as therapeutic function, immunogenicity, cancer cell specificity etc (*9*). Numerous clinical trials of a variety of immune-based therapies targeting the tumor-associated antigen MUC1 have been reported, including MUC1 vaccines, anti-MUC1 antibodies and T cells engineered with chimeric antigen receptors. Chemically, MUC1 is a type I transmembrane protein with a heavily glycosylated extracellular domain composed of 20-120 tandem repeats of twenty amino acids (GSTAPPAHGVTSAPDTRPAP), each with five potential *O*-glycosylation sites at Ser and Thr residues (*10, 11*). Its *O*-glycosylation is initiated by UDP-GalNAc:polypeptide *N*-acetylgalactosaminyltransferases (*12*), and further elongated by Core 1 and branched Core 2 enzymes, protecting the underlying epithelia from desiccation and pathogen invasion. In most human epithelial cancers, MUC1 is overexpressed and loses its polarity. Especially, MUC1 in tumor cells is aberrantly glycosylated which is caused by genetic deficiency of COSMC (core-1 β3-Gal-T-specific molecular chaperone (*13, 14*) which forms complex with core-1 β3-Gal-T), resulting in termination of glycosylation and exposure of immunogenic truncated glycans including Tn-antigen (GalNAc-*O*-S/T) and sialyl-Tn-antigen (Neu5Acα2-6GalNAc-*O*-S/T). Monoclonal antibodies binding to consecutive GalNAc residues located on MUC1, the MUC1 peptide sequence, or both GalNAc and peptide sequence have been reported (*8*). COSMC deficiency in mice causes spontaneous autoimmunity, suggesting the role of protein *O*-glycosylation in homeostasis of immune defense and immune tolerance (*15*).

While aberrantly glycosylated MUC1 is a viable anticancer immunotherapy target, vaccine strategies incorporating unglycosylated MUC1 fragments have been largely unsuccessful, probably due to the conformational discrepancies between the unglycosylated MUC1 peptide vaccine and aberrantly glycosylated tumor-associated MUC1 (*16, 17*). Thus, the incorporation of MUC1 glycopeptide vaccine candidates has been new focus of research (*6*). The knowledge regarding the rule that govern the immunogenicity of certain MUC1 glycopeptide are still scarce. Moreover, despite a plethora of anti-MUC1 mAbs having been developed, both the molecular mechanism by which MUC1 specific antibodies recognize their target and the role of glycosylation within the epitope are not well understood. Crystallography along with computational approach will help decipher the antibody recognition of glycan epitope and provide structural insights on epitope selection for vaccine design. To date, only three crystal structures of Muc1-specific antibodies in complex with their peptide and corresponding Tn-glycopeptide epitope were reported respectively: mAb SM3 (*18*), which was raised with partly deglycosylated mucin from human milk; mAb AR20.5 (*19*), which was raised by immunization with MUC1 from an ovarian cancer patient; and SN-101, which was raised by immunization with synthetic MUC1 glycopeptide (*20*). All the antigen epitopes of SM3, AR 20.5 and SN-101 include the immunodominant PDT(GalNAc) region of MUC1 fragments, while SN-101 binds to additional non-linear amino acid residues located in its conformational epitope.

To understand the interactions that govern recognition by tumor-derived MUC1 by anti-MUC1 antibodies at the molecular level to guide structure-aided vaccine design, we selected two individual linear glycopeptide-specific anti-MUC1 antibodies, 14A and 16A (*21*), for structural study. 14A and 16A, were raised by immunizing mice with MUC1-transfected mutant cancer cells deficient of mucin-type core-1 β1-3 galactosyltransferase activity and screening against chemically synthesized glycopeptide RPAPGS(Tn)TAPPAHG. Several crystal structures of two glycopeptide-specific antibody fragments (Fabs) of 14A and 16A in complexed with the antigen peptide, S-modified monoglycopeptide, T-modified monoglycopeptide and ST-diglycopeptide were determined at atomic resolution. These structures showed that both antibodies recognized a novel epitope within the repeat region distinct from previously reported immunodominant PDTR motif. Combined with detailed mutation analysis and molecular dynamics simulation, we evaluated how the Tn-glycosylation at different site of MUC1 peptide backbone influences the recognition between antigen and antibody. Structure elucidation to understand and rationalize how glycosylation can modify the structure of a cancer antigen, and how these modifications can affect the tumor-associated-antigen targeted antibodies’ binding affinities, will be important for the design of MUC1-based vaccines with improved immune response and the development of high-affinity therapeutic anti-MUC1 antibodies.

## Results

### COSMC deficiency caused increased binding to 14A and 16A mAbs

We compared the binding affinity of 16A and 14A mAbs to MUC1 protein expressed by 293T-COSMC^-/-^ cell line which are depleted of COSMC gene through CRISPR-Cas9 technology, and its parent cell line. Flow cytometry analysis showed that COSMC deficiency caused significantly increased binding affinity to 16A, but not 14A (Fig. 1), as shown by increased staining signal measured by flow cytometry. Western blot analysis confirmed the binding of 16A and 14A are specific to COSMC^-/-^ cell surface expressed MUC1 protein, but not on the normal HEK293 cells (Fig. 1). The cell surface expressed MUC1 might represent more natural conformations for these exposed tandem repeat with truncated hypoglycosylation only revealed by flow cytometry staining.

**Fig. 1.**
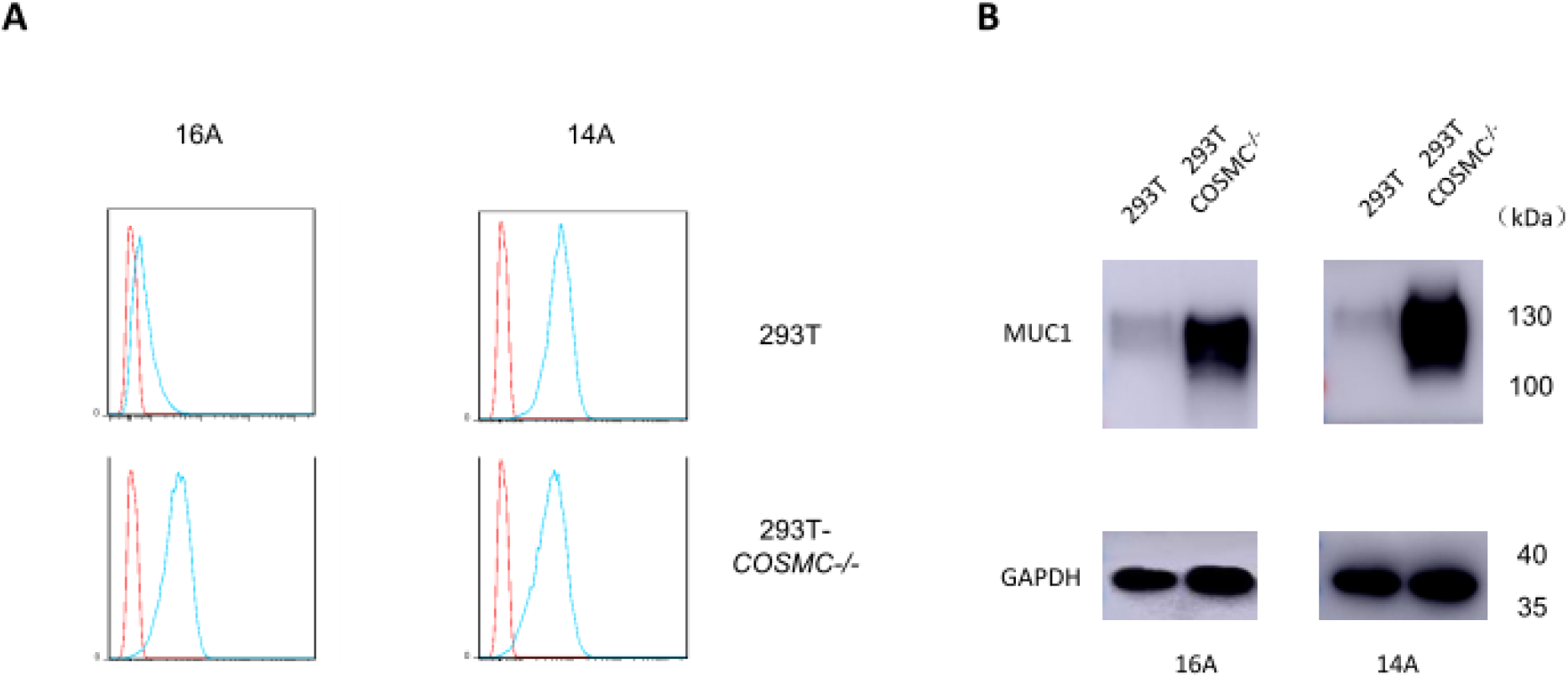
Differential binding to COSMC^-/-^ cells by 16A mAb. (**A**) Staining of 293T and 293T-COSMC^-/-^ cells by 14A and 16A antibodies (mouse IgG1, 0.2 μg/ml) followed by fluorescence-labelled secondary antibodies. (**B**) Western blot analysis of protein lysates from 293T and 293T-COSMC^-/-^ cells.

### Epitope Mapping 14A and 16A mAbs

By glycopeptide array using 73 glycopeptides containing different regions of MUC1 tandem repeats (table S1), we clearly identified GST(T_N_)A region as the epitope for both 16A and 14A mAbs (Fig. 2). Negative binding to glycopeptides 1-12 and 48-59 excluded the possibility of recognition to PDTR or GVTS regions by 14A and 16A. The strongest binding was found for glycopeptide 35, which contained two tandem repeat of GST(T_N_)A sequences, suggesting simultaneous bivalent-recognition of the two epitopes on the glycopeptide by the two Fabs of the antibodies. This is consistent with the observation that glycopeptide 13, which only contained one GST(T_N_)A repeat, also showed more than 50% reduced binding signals. Glycosylation of the peptide seemed to enhance the binding of the antibodies. A non-glycosylated GSTA epitope (glycopeptide 15, 16, and 32) reduced the binding affinities by more than 50% compared to the glycosylated counterpart (glycopeptide 35).

**Fig. 2.**
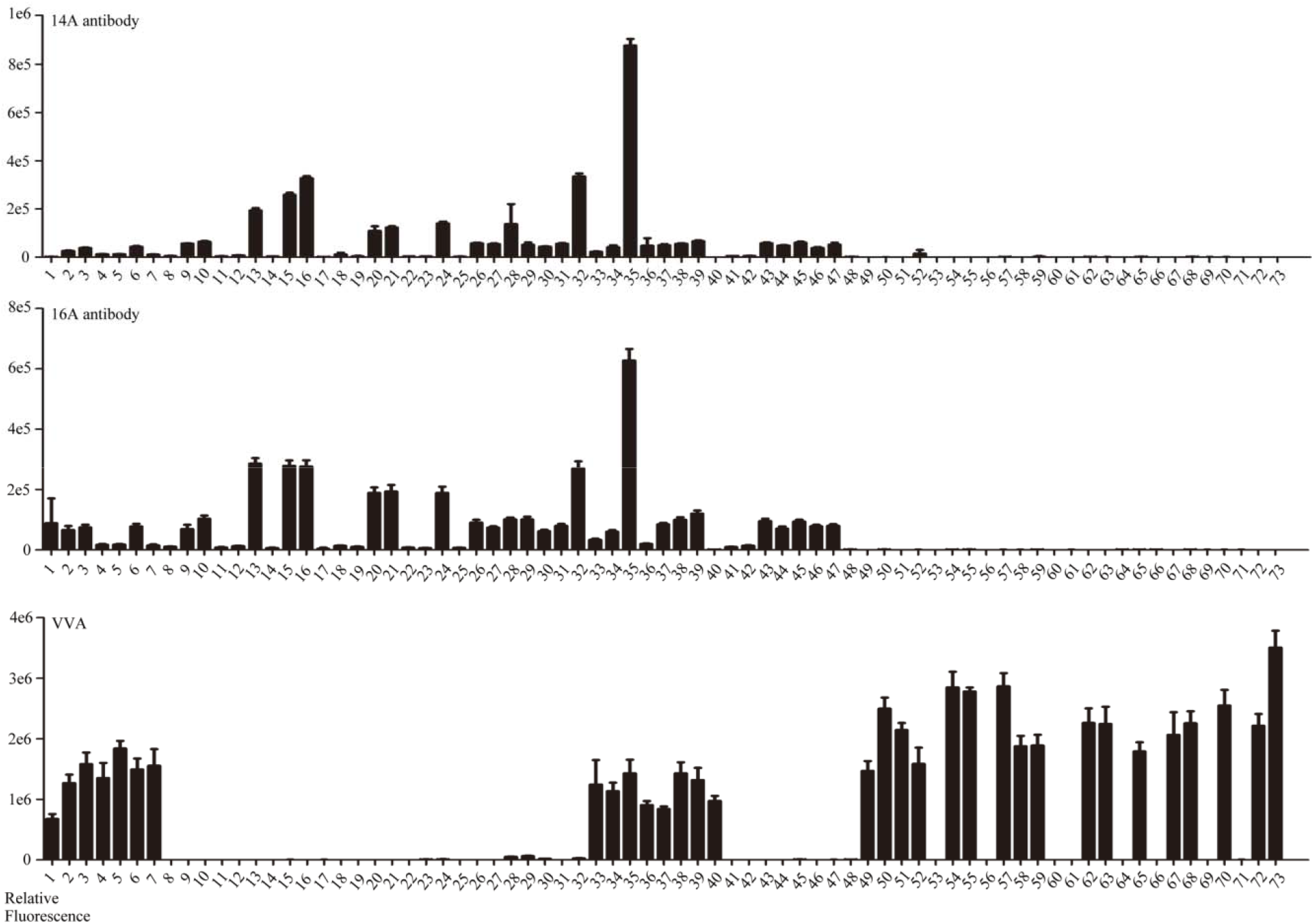
Glycopeptide array of 14A and 16A antibodies. Antibodies and Vicia Villosa Lectin (VVA) control were incubated with 73 glycopeptides containing different regions of MUC1 tandem repeats (table S1).

### Affinity and Specificity of Fab Fragments of 14A and 16A mAbs

Both antibodies, 14A and 16A mAbs, were initially identified by screening against synthesized glycopeptide RPAPGS(Tn)TAPPAHG. To decipher the impact of glycosylation of the epitopes on 14A and 16A binding affinity to MUC1, a RPAPGSTAPPAHG as glycopeptide backbone was used. The peptide backbone contains two putative glycosylation sites at Ser and Thr, thus, 4 (glyco)peptide were synthesized and tested. These include a non-glycosylated peptide (RPAPGSTAPPAHG), glycopeptides bearing a single GalNAc on Ser (RPAPGS(Tn)TAPPAHG, GalNAc-S) and on Thr (RPAPGST(Tn)APPAHG, GalNAc-T), and glycopeptide bearing two GalNAc on Ser and Thr (RPAPGS(Tn)T(Tn)APPAHG, GalNAc-ST) (Fig. 3A). Surface plasmon resonance was used to assess the binding specificities and affinities of the MUC1-specific mAbs for these (glyco)peptide antigens (Fig. 3B). Specifically, 14A bound to the non-glycosylated peptide with a KD of 114.6 nM, while the value for 16A is 220 nM (Table 1). Addition of a single GalNAc on Thr dramatically enhanced the binding of antigen both for 14A and 16A, with the KD decrease 11.6-fold (9.9 nM) and 30.6-fold (7.2 nM), respectively, compared to the non-glycosylated peptide. In contrast, adding a single GalNAc on Ser only slightly enhanced the binding of 14A at 1.6-fold (73.5 nM), whereas it has almost no effect on 16A (219.5 nM). The two Tn modified glycopeptides at both Thr and Ser position bound to 14A with a KD of 1.4 nM, about 52.5-fold binding enhancement compared with the GalNAc-S, and 81.9-fold binding enhancement compared with the non-glycosylated peptide, indicated that the effect of dual-glycosylation on Ser and Thr is additive for 14A. 16A bound to GalNAc-ST with a KD of 6.7 nM, which is almost identical with the GalNAc-T, further confirmed that GalNAc addition on Ser has no effect for 16A binding.

**Fig. 3.**
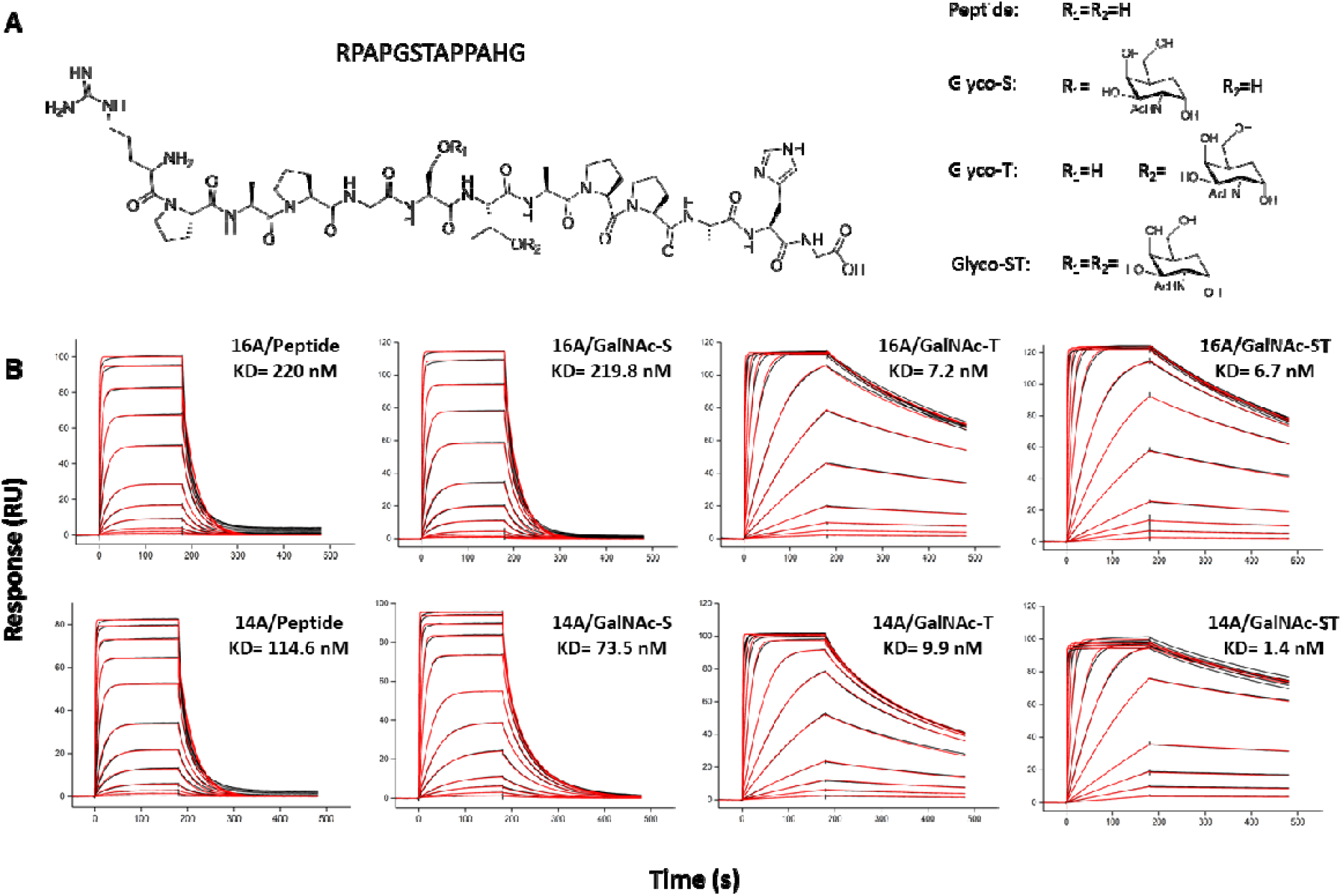
Affinity and Specificity of Fab Fragments of 14A and 16A mAbs. (**A**) Chemical structure of synthesized (glyco)peptide used for SPR analysis and crystallization. (**B**) SPR sensorgram overlays of (glyco)peptide (concentrations of 1, 2.5, 5, 10, 25, 50, 100, 250, 500, 1,000, 2,500 and 5,000 nM) binding to 16A and 14A Fab. Black lines indicate observed data points; red lines indicate fitted data.

**Table 1.** SPR measurement of dissociation constants for the binding of 14A and 16A Fabs to (glyco)peptides.

| (Glyco) peptide | ka (1/Ms) | kd (1/s) | KD (nM) | Rmax (RU) |
| --- | --- | --- | --- | --- |
| **16A WT** |  |  |  |  |
| Peptide | 153494.6 | 0.033787 | 220.0 | 86.6 |
| Glycopeptide-S | 150642.8 | 0.033116 | 219.8 | 105.2 |
| Glycopeptide-T | 333366.1 | 0.002392 | 7.2 | 114.9 |
| Glycopeptide-ST | 289028.4 | 0.001937 | 6.7 | 124.8 |
| **16 W34F** |  |  |  |  |
| Peptide | 86160.0 | 0.295400 | 3429.0 | 59.6 |
| Glycopeptide-S | 86210.0 | 0.336800 | 3907.0 | 85.1 |
| Glycopeptide-T | 73610.0 | 0.089950 | 1222.0 | 112.4 |
| Glycopeptide-ST | 79700.0 | 0.090090 | 1130.0 | 127.6 |
| **16 W34A** |  |  |  |  |
| Peptide | — | — | — | — |
| Glycopeptide-S | — | — | — | — |
| Glycopeptide-T | — | — | — | — |
| Glycopeptide-ST | — | — | — | — |
| **16 R32G** |  |  |  |  |
| Peptide | 162951.3 | 0.053169 | 326.3 | 79.4 |
| Glycopeptide-S | 174472.5 | 0.029398 | 168.5 | 94.4 |
| Glycopeptide-T | 359105.5 | 0.008231 | 22.9 | 108.1 |
| Glycopeptide-ST | 533726.7 | 0.001825 | 3.4 | 104.8 |
| **14A WT** |  |  |  |  |
| Peptide | 268707.5 | 0.030806 | 114.6 | 68.0 |
| Glycopeptide-S | 640975.0 | 0.047107 | 73.5 | 94.0 |
| Glycopeptide-T | 521101.4 | 0.005157 | 9.9 | 99.3 |
| Glycopeptide-ST | 928676.0 | 0.001298 | 1.4 | 98.0 |
| **14 G32R** |  |  |  |  |
| Peptide | 191198.1 | 0.023278 | 121.7 | 74.2 |
| Glycopeptide-S | 179521.0 | 0.022909 | 127.6 | 86.6 |
| Glycopeptide-T | 313691.6 | 0.001590 | 5.1 | 91.9 |
| Glycopeptide-ST | 302506.7 | 0.001371 | 4.5 | 106.1 |

### Crystal Structures of Fab 16A in complex with antigen peptide and glycopeptide

To ascertain the structural basis for the effect of antigen peptide glycosylation on antibody-antigen binding, we screened the co-crystallization conditions for 16A Fab with the synthetic MUC1 peptide and glycopeptide. The 16A/Peptide, 16A/Glyco-T, and 16A/Glyco-ST complexes were crystallized in the space groups P12_1_1, P2_1_2_1_2_1_, and P2_1_2_1_2_1_, respectively, and the structures were solved and refined to 2.10 Å, 1.56 Å, and 2.20 Å resolution, respectively (table S2). In all three complex structures, their asymmetric units contain only one 16A Fab (Fig. 4A). The overall conformation of the Fab backbone was nearly identical in all (Glyco)peptide complexed structures of 16A Fab.

**Fig. 4.**
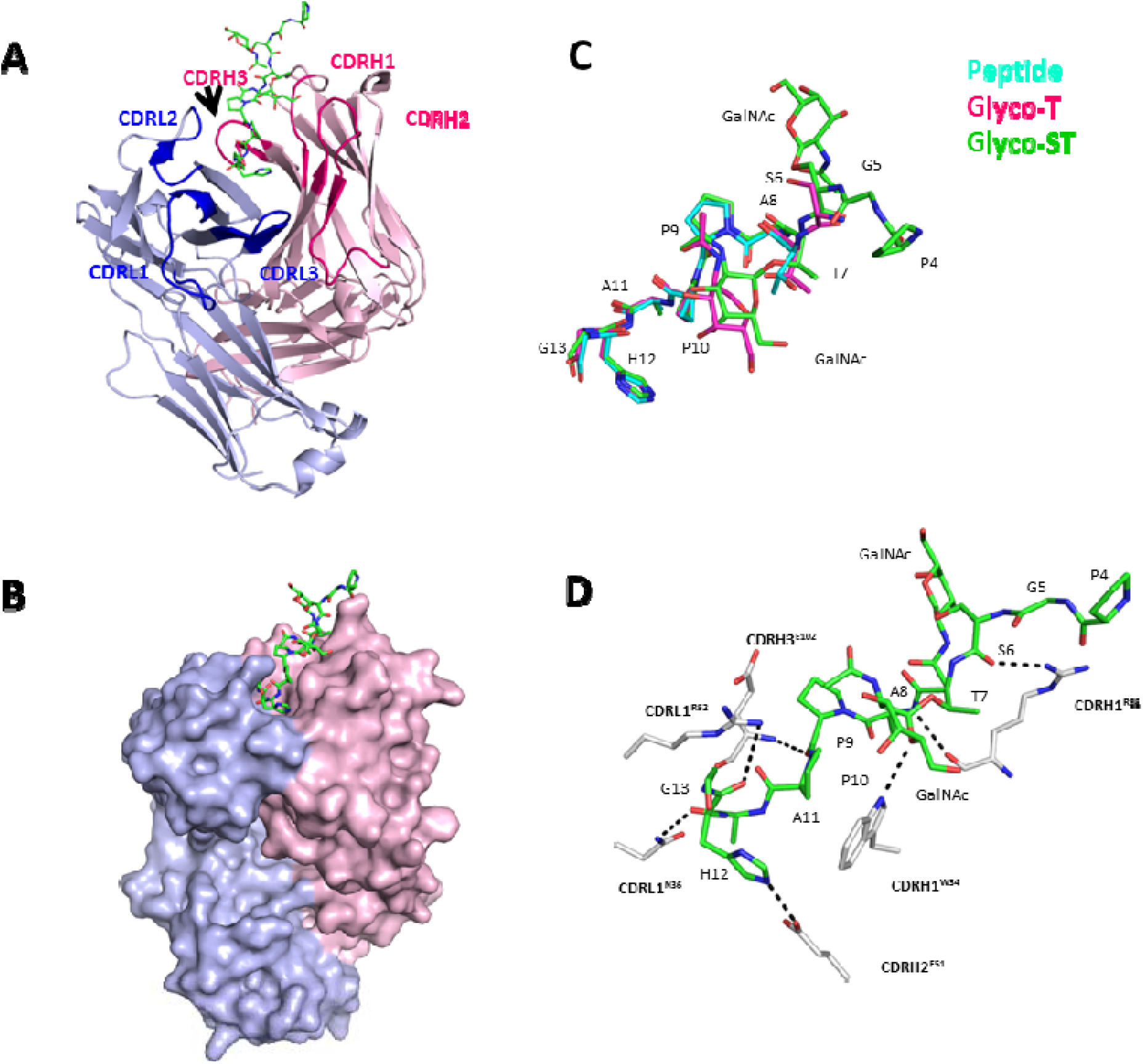
Structural characterizations of the 16A Fab/(Glyco)peptide complexes. (**A**) The overall structure of the 16A Fab/Glyco-ST complex, with the Glyco-ST carbon atoms colored in green, and L and H chain colored in light blue and light pink, respectively. The three light-chain and three heavy-chain CDRs of 16A that are involved in binding (glyco)peptide are highlighted in blue and hot pink, respectively. (**B**) The surface structure of the 16A Fab/Glyco-ST complex. (**C**) Superposition of the peptide backbone of compounds peptide, Glyco-T, and Glyco-ST in complex with 16A. (D) Detailed interactions between 16A Fab and Glyco-ST. Dashed lines represent hydrogen bonds. Epitope of peptide sequence is RPARPAPGSTAPPAHG

The Fabs were numbered with the Kabat system, and the peptides were numbered from 1 to 13, respectively. In the structure of 16A in complex with a synthetic MUC1 peptide (RPAPGSTAPPAHG), the region corresponding to TAPPAHG exhibited clear electron density in 2*F*_*o*_-*F*_*c*_ maps. The six N-terminal amino acids of the synthetic peptide (RPAPGS) were disordered according to the electron density maps, suggesting that these residues do not form stable interactions with the 16A Fab. The 16A paratope forms a surface groove using CDRs L1, L2, L3, H1, H2 and H3 (Fig. 4B). Surprisingly, the MUC1 peptide epitope makes only a few polar contacts with the paratope, wherein the light chain of 16A forms a total of two hydrogen bonds to the epitope (CDRL1^N36^ to A11, CDRL1^R52^ to H12) and the heavy chain forms three hydrogen bonds to the ligand (CDRH1^R32^ to A8, CDRH2^E51^ to H12 and CDRH3^E102^ to P9).

The structure of 16A in complex with a GalNAc glycosylated glycopepitde on Thr (RPAPGST(Tn)APPAHG, denoted as Glyco-T) revealed unambiguous electron density in 2*F*_*o*_-*F*_*c*_ maps for the segment of ST(GalNAc)APPAHG and the carbohydrate portion on Thr. Unlike the non-glycosylated peptide structure, the Ser (S6) was visible in this complex, albeit with a high B-factor. The glycopeptide lies in the same surface groove as the non-glycosylated peptide (Fig. 4C). The interactions of 16A with the peptide portion of glycopeptide were essentially identical to those observed in the peptide structure, except an extra H-bond between CDRH1^R32^ and S6. Although the GalNAc residue points away from the binding site toward the solvent, an hydrogen bond exists between the endocyclic oxygen O5 of GalNAc and the NH group of CDRH1^W34^ of 16A (Fig. 4D). Moreover, the sugar ring stacks with the aromatic ring of CDRH1^W34^, further providing the impetus for the observed selectivity of 16A for GalNAc-containing antigens.

The structure of 16A in complex with glycopeptide bearing GalNAc on both Ser and Thr (RPAPGS(Tn)T(Tn)APPAHG, GalNAc-ST) revealed clear electron density for the region corresponding to the PGS(GalNAc)T(GalNAc)APPAHG and the carbohydrate portion of the antigen. The interactions of 16A with the GalNAc-ST were the same to those observed in the Glyco-T structure. The GalNAc carbohydrate on Ser does not appear to directly participate in the binding.

### Crystal Structures of Fab 14A in complex with antigen glycopeptide

To investigate the structural basis of binding preference of 14A toward glycopeptide bearing GalNAc glycosylation at Ser or Thr, and to decipher the differences between 14A and 16A on their recognition of COSMC^-/-^ cell surface expressed MUC1, 14A Fab was co-crystallized with each of its antigen glycopeptide (Glyco-S, Glyco-T, and Glyco-ST) in the space groups P12_1_1, P2_1_2_1_2_1_, and P2_1_2_1_2_1_, and the structures were refined to 2.73 Å, 3.20 Å, and 3.50 Å, respectively (table S2). All structures contained three Fabs arranging in a head-to-tail fashion per asymmetric unit (Fig. 5A). The overall conformation of the Fab polypeptide backbone was nearly identical in all the glycopeptide complex structures of 14A Fab and is similar to that of the 16A Fab (0.21 Å root-mean-square deviation (rmsd) between 14A/Glyco-ST and 16A/GlycoST). Clear electron density was evident for residues PGSTAPPAHG of Glyco-S, and for residues GSTAPPAHG of Glyco-T and Glyco-ST glycopeptides in the 2*F*_*o*_-*F*_*c*_ maps. The carbohydrate portion of the three glycopeptides antigen also exhibit well-defined electron density in 2*F*_*o*_-*F*_*c*_ maps. Moreover, the overall conformation of the peptide fragment of the different MUC1 glycopeptide is nearly identical to that found in the crystal structure of 16A complex (Fig. 5C).

**Fig. 5.**
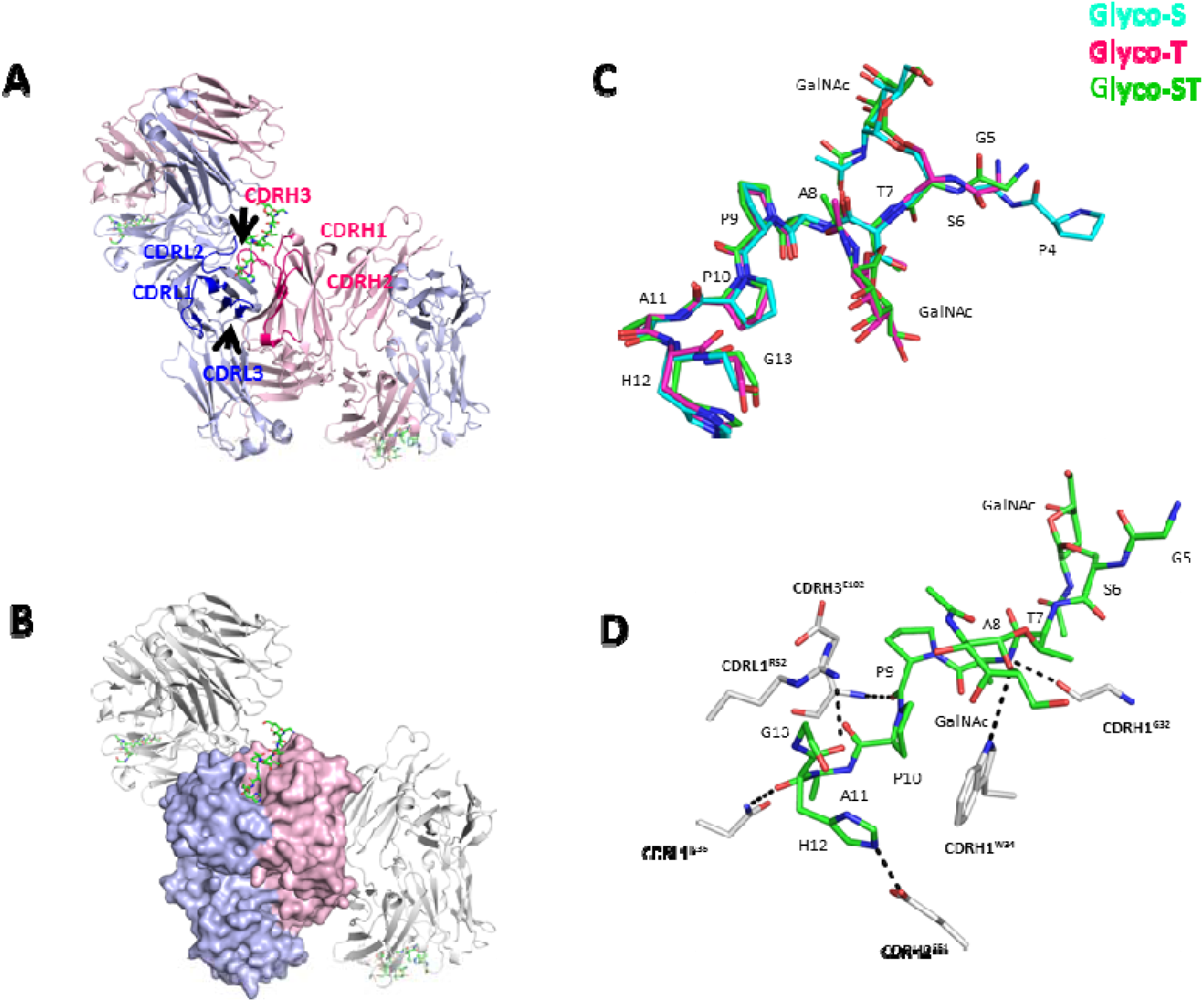
Structural characterizations of the 14A Fab/(Glyco)peptide complexes. (**A**) The overall structure of the 14A Fab/Glyco-ST complex, with the Glyco-ST carbon atoms colored in green, and L and H chain colored in light blue and light pink, respectively. The three light-chain and three heavy-chain CDRs of 14A that are involved in binding (glyco)peptide are highlighted in blue and hot pink, respectively. (**B**) The surface structure of the 14A Fab/Glyco-ST complex. (**C**) Superposition of the peptide backbone of compounds Glyco-S, Glyco-T, and Glyco-ST in complex with 14A. (**D**) Detailed interactions between 14A Fab and Glyco-ST. Dashed lines represent hydrogen bonds. Epitope of peptide sequence is RPARPAPGSTAPPAHG.

The glycopeptide antigen lies within a surface groove formed by CDRs L1, L2, L3, H1, H2 and H3 of 14A (Fig. 5B). Close inspection of the ligand-binding pocket revealed that the peptide portion of glycopeptide makes only a few polar contacts with the antibody-combining site. The light chain of 14A forms a total of two hydrogen bonds to the peptide epitope (CDRL1^N36^ to A11, CDRL1^R52^ to H12) and the heavy chain forms three hydrogen bonds to the peptide portion of glycopeptide, (CDRH1^G32^ to A8, CDRH2^E51^ to H12, CDRH3E^102^ to P9) (Fig. 5D). In the 14A/Glyco-S structure, the GalNAc does not appear to directly participate in the interaction, as it makes no specific polar contacts to the antibody paratopes. Therefore, the enhanced binding between 14A and GlycoS cannot be explained based on the complex structure. On the other hand, in the structures of 14A/Glyco-T and 14A/Glyco-ST, an intermolecular hydrogen bond exists between the endocyclic oxygen O5 of GalNAc on Thr site and the NH group of CDRH1^W34^ of 14A (Fig. 5D), as well as sugar ring stacking interaction with the aromatic ring of CDRH1^W34^. Moreover, the GalNAc residue on GalNAc-T forms intramolecular hydrogen bonds between its C3 OH and Ac oxygen group. These observations shows structural basis for enhanced binding for Glyco-T and Glyco-ST.

### Mutation analysis

The structural analysis of 14A and 16A in complex with their (glyco)peptides have unveiled the potential antigen-antibody interaction contributing to enhanced binding affinity to the glycopeptide in detail. To further assess the roles of residues pertaining to the binding specificity, a series of Fab mutants were constructed, and their binding affinities toward four (glyco)peptides were examined by SPR (Table 1).

Both the structures of 14A and 16A in complex with GalNAc-T suggested that the observed higher affinity to the Thr-glycosylated peptide for 14A and 16A was mediated by a hydrogen bond between the endocyclic oxygen O5 of GalNAc and the NH group of CDRH1^W34^ indole ring and stacking of the sugar ring to the aromatic ring of CDRH1^W34^. We choose 16A as the model system to study the effect of CDRH1^W34^. First, we mutated the W34 to Alanine to eliminate both the hydrogen bond and stacking interaction. The results show that this mutant completely lost its binding ability to all four (glyco)peptides. Then we measured the binding of W34F mutant, which disrupted the hydrogen bond interactions, but preserved the π-π stacking interaction. Although W34F mutation resulted in decreased binding affinities for all test (glyco)peptides, the preference for GalNAc-T was also weakened compared with the non-glycosylated peptide, with the ratio of KD_peptide_/KD_GalNAc-T_ reduced from 30.6-to 2.8-fold. Interestingly, the loss of affinity preference of W34F appeared to be primarily due to the disappearance in the dissociation rate differences (off rate; *k*_d_) for peptide and GalNAc-T, while differences in dissociation rate mostly conferred the binding differences of the wild-type 16A towards two peptides.

Sequence alignment (fig. S3) and structure analysis indicated that only residue 32 in heavy chain is different in the antigen binding site between 14A (CDRH1^G32^) and 16A (CDRH1^R32^), suggesting it may be related to the observed affinity differences towards Ser-glycosylated peptide. Moreover, Comparison of the structure complexes of 16A:GalNAc-T and 16A:GalNAc-ST showed that there is a sidechain movement for CDRH1^R32^ when 16A binds glycopeptide bearing GalNAc modification on Ser6. Therefore, we carried out studies to examine the role of residue 32 in the GalNAc-S recognition. We introduced a glycine at CDRH1^R32^ to glycine in 16A, while mutated CDRH1^G32^ to arginine in 14A. As shown in Table 1, when CDRH1^R32^ was mutated to glycine, the KD_peptide_/KD_GalNAc-S_ was ∼1.9-fold, indicating 16A obtained affinity preference toward Ser modified glycopeptide. In contrast, the G32R of 14A showed almost identical KD for both peptide and GalNAc-S (121.7 and 127.6 nM, respectively), indicating 14A lost its preference toward GalNAc-S. These results demonstrate the important role of residue 32H in GalNAc-S binding.

### Simulations revealing dynamical interactions between MUC1-peptides and antibodies

Based on crystal structures, we constructed systems with 10-residue protopeptides similar to the MUC1 peptides for molecular dynamics simulations (without N-terminal RPA residues, see Method section for details). Simulations revealed that the peptide forms stable interactions with antibodies via the segment of Thr4 to His9, indicated by the low conformational fluctuations in all simulated systems (see fig. S1a). The residues near the peptide termini exhibited larger fluctuations, indicating more flexible conformations. This is consistent with the crystallography data, from which the electron density of N-terminal residues are not well resolved.

We extracted detailed hydrogen bonding information between the peptides and the antibodies from simulation trajectories (table S1). The most frequently observed hydrogen bonds (with occupancy ∼80%) are Pro6:Glu102 and Ala5:Gly32(14A) or Ala5:Arg32(16A), where the residues of the MUC1 peptide and of the antibodies (heavy chain) are shown on the left and right of the colon respectively. The Asn36 and Arg52 of the light chain antibody also form noticeable hydrogen bonding interactions with peptides via Ala8 and His9 respectively. Besides the hydrogen bonding interactions between the peptide and the antibodies, simulations show that hydrophobic interactions contribute to the peptide-antibody recognition. For example, the Ala5 of MUC1-peptide and the Tyr33 of the antibody heavy chain are in close contact throughout the simulations, reflected on the distance between the Ala5 of the MUC1 peptide and the Tyr33 benzene ring (fig. S2). Similarly, we identified the hydrophobic contacts between Ala8 of the MUC1-peptide and the Tyr100 of the heavy chain. These key interactions are summarized in fig. S1b for both 14A and 16A antibodies.

According to SPR data, the wild type 14A Fab shows better affinity to MUC1-peptides than 16a for almost all glycosylation conditions (see Table 1). Interestingly, the mutation of Gly32 to Arg32 in heavy chain of 14A_G32R led to a decreased association constants (Ka) compared to the wild type 14A Fab, while the 16A_R32G and the mutation of Arg32 to Gly32 in heavy chain of 16A Fab led to larger Ka. Based on the crystal structures and simulation trajectories, we found that the residue on site 32 of heavy chain may affect the binding via the following interactions: (1) the direct interaction between the side chain of residue 32 and the peptide; (2) the indirect influence by the conformation of Tyr100, which is related to the peptide binding. As revealed from the hydrogen bond analysis, the Gly32 or Arg32 are crucial for peptide binding, and the Arg32 undergoes larger fluctuations due to its longer side chain. The dynamical feature is reflected on the distance between the C_Z_ atoms of R32 and Y33. In the apo form of 16A, this distance has a broader distribution, between 3 Å and 13 Å; with MUC1-peptide bound to 16A, this distance is narrowed to a range between 5 Å and 10 Å. We also observed that the Tyr100 adapts a specific conformation for the peptide binding. As shown in fig. S1c, the Tyr100 side chain orientations are different in the apo and MUC1-peptide bound forms of 16A. Similarly, the G32R of 14A Fab shows a bended conformation of Tyr100 that is similar to the wild type 16A (fig. S1b). This is consistent with the reduced Ka in G32R of 14A Fab and the increased Ka in R32G of 16A Fab.

Simulations revealed that the glycosylation of Ser3 and Thr4 in MUC1-peptide could introduce extra hydrogen bonds with antibodies. The glycosyl groups linked on Thr4 interact with Trp34 and Asp55, in heavy chain of both 14A and 16A Fabs, providing structural basis for the stronger binding affinity between glycosylated peptide (Glyco-T) and the Fabs, as compared to the non-glycosylated peptide (Table 1). Although the Glyco-S formed new hydrogen bonds with Asp29 and Tyr101 in heavy chains, enhancements of Fab binding were not observed in the SPR analysis, suggesting that Ser3 is not in the core region of Fab binding.

## Discussion

### Paratopes that discriminate normal and tumor glycans

Several antibodies that target abnormally glycosylated proteins in cancer have been co-crystalized with their ligand counterpart. One class of antibodies bind to exposed Tn sugar in a “pocket” mode (*22*–*26*), as represented by 237 mAb which recognizes a glycopeptide from podoplanin (*22, 23*). From the complexed structure between 237 mAb and the gloycopeptide, the Tn sugar buried in a deep pocket, while the peptide moiety binds in a shallow grooved formed by extensive somatic mutations in the framework region. Both the sugar and peptide moieties’ interaction mode confer 237mAb exquisite selectivity toward Tn-antigen, while the antibody shows lower specificity to peptide portion (*23*). The heavy chains of this class of antibodies include the usage of several VH genes with certain degree of sequence identities, according to our studies on mAb clones binding to clustered Tn antigens conjugated to Bovine serum albumin (Table 2). The “pocket” binding to Tn antigen does not allow binding to further elongated glycans, which is the basis for binding to COSMC-deficient cells. The other class of mAbs have been reported to be specific to peptide backbone of glycopeptides, represented by the PDT(GalNAc)R region of MUC1. For example, mAb SM3 was reported as binding to both sugar and peptide portion of PDT(GalNAc)RP sequence. Nevertheless, the presence of sugar moiety of the glycopeptide does not provide additional interactions with the paratope of the antibody (*18*). Similarly, mAb AR20.5 was found to bind to peptide portion of DT(GalNAc)RAP sequence, while direct interaction between antibody and sugar portion were not observed. It is hypothesized that the sugar portion regulates the intrinsic flexibility of the glycopeptide, thus could decrease the entropy loss while interaction with the antibody, thus increasing its affinity in antibody binding (*19*). SN-101 mAb was reported as binding to both GalNAc sugar and the conformational peptide residues including PDT region and other non-linear amino acid residues (*20*), while the GalNAc attached to PDT region was found to be critical for effective binding. The affinity of SN-101 mAb’s binding to MUC1 glycopeptide is relatively low (KD=0.5 μM), as the glycosylation of other two amino acids within the conformational epitope actually blocks the SN-101 binding (*20*).

**Table 2.** IGHV gene families of mAbs binding to Tn antigen without peptide specificity.

| mAb | IGHV gene | Isotype | Specificity | Reference |
| --- | --- | --- | --- | --- |
| 17B | IGHV1-78*01 | IgM | BSA-conjugated Tn | this study |
| 54B | IGHV1-72*01 | IgM | BSA-conjugated Tn |  |
| 60A | IGHV1-34*01 | IgM | BSA-conjugated Tn |  |
| 74A | IGHV5-6*01 | IgM | BSA-conjugated Tn |  |
| Kt-IgM-8 | IGHV9-3 | IgM | Tumor-expressed Tn | 24 |
| 83D4 | IGHV1S53 | IgM | Tumor-expressed Tn | 25 |
| 237 | IGHV1S53 | IgG2a | Tumor-expressed Tn | 22-23 |
| B72.3 | IGHV1S53 | IgG1 | Tumor-expressed Tn | 26 |

In contrast to above-mentioned mAbs, we have found clear interaction between the GalNAc sugar residue of S(Tn)T(Tn)APPAHG sequence and the CDRH3 region in the 14A and 16A Fabs characterized in this study. Furthermore, molecular simulation studies suggest that the GalNAc residue of S(Tn)T(Tn)APPAHG also significantly alters the flexibility of the peptide, and generates a favorable confirmation for antibody binding. In a western blot analysis for 14A and 16A recognition toward MUC1 protein expressed in COSMC-deficient cells, they possess comparable recognition specificities, only recognized Tn-MUC1 protein, instead of normal MUC1 with full O-glycosylation chain, which has been suggested to prevent the accessibility to MUC1 peptide backbone. With denatured condition, the fully glycosylated MUC1 peptide backbone expressed in normal cells could be even more occluded from accessibility from the antibody binding. By flow cytometry measurement to cell surface binding, 16A but not 14A could distinguish between the tumor cell surface expressed hypoglycosylated MUC1 and the MUC1 expressed by normal cells. The reason could be that even though MUC1 expressed by the normal cells and tumors have distinguished glycosylation, 14A’s epitope binding pocket seems to not evolve as advanced as 16A to distinguish between normal elongated glycosylation chain vs. truncated glycosylation. Recognition pattern for glycopeptide is both protein sequence, glycosylation on the tandem repeat position and glycosylation type dependent.

From complexed structures of 16A interacted with Tn-glycopeptides, 16A seems to achieve better recognition specificity toward more truncated glycosylated-MUC1 specifies by steric hinderance imposed by the key residues aligned at the entrance of the binding pocket, especially Arg 32, a corresponding Gly for 14A. There are extra hypermutations occurred at the frame region of 16A; but not in 14A. FWR mutations have been demonstrated to be selected during early development of bnAb and alter the Fab elbow angle with increased interdomain flexibility; sometimes associated with enhanced neutralization of broadly neutralizing antibody (*27*). Whether the somatic mutations at 16A FWR also enable 16A to adopt the thermal stability and flexibility and fine selectivity towards more truncated sugar-substituted epitopes worth further studies.

### Immunogenicity of peptide region modified by consecutive glycosylation sites

Dense glycosylation of peptide region is the main barrier when such linear glycopeptides are tested as vaccine candidates. Our results showed the generation of high-affinity antibodies toward linear peptide region, with increased affinity to glycan-modified peptide (KD=6.7 nM). The immunogenicity of STAPPAG region might be associated to its unique secondary structure. Previous NMR studies on this region suggested its non-globular nature. Both peptide and carbohydrate contributed to a folded structure in solution (*16*). In another study by NMR on glycosylated GSTAPPAHGVTSAPDTRPAP, glycosylation at GVTS region formed an extended, rod-like secondary structure. The APDTR region formed a turn structure which is more flexibly organized (*17*). We have generated monoclonal antibodies specific to either GVTS or PDTR glycopeptides (data not shown) with enhanced binding to glycosylated linear peptide regions. Further structural analysis will help to elucidate the immunogenicity of this highly glycosylated tandem repeating units.

### Paratopes selectively recognize the proximal site of glycosylation of bound peptide by peptide-binding CDR groove with a glycan length-limiting edge

Our study demonstrated a paratope-binding mode where the proximal site of glycosylation significantly increases the affinity of binding to bound peptide. This finding may have implications for antibody binding to other peptide region modified by consecutive glycosylation sites. We have also isolated another monoclonal antibody, TJ7, binding to PAHGVT(Tn)SAPD but not PAHGVT(Tn)S(Tn)APD (unpublished data).

Our finding may also help to systemically identify immunologic self-peptide epitopes in autoimmune diseases. For example, the IgA1 hinge region, PVPSTPPTPSPSTPPTPSPS, is a target for antibody binding when the O-glycosylation is blocked and Tn antigen (GalNAc) is exposed (*13*). This region contains 6 O-glycosites, while which GalNAc residues are involved in disease pathogenesis remain to be elucidated. Similarly, the nature of paratopes that bind to GalNAc modified proteins are unknown in Tn polyagglutination syndrome, a clonal disorder characterized by the polyagglutination of red blood cells by naturally occurring anti-Tn antibodies following exposure of the Tn antigen on the surface of erythrocytes. Identifying the paratopes of antibodies reactive to GalNAc-modified proteins may provide clues to earlier diagnosis and treatment.

## Materials and Methods

### Measurement of mAb binding to cells by flow cytometry

HEK293T-cosmc cell line which is genetically depleted of COSMC was generated by CRISPR-cas9 knockdown method (details will be published elsewhere). 2×10 ^5^ HEK293T-cosmc cells and wild type HEK293T cells were incubated with various concentrations of mAbs 16A/14A in 50 μl of phosphate buffer (pH7.4) containing 1% BSA and 2 mM EDTA for 30min at 4°C. Cells were washed three times in PBS buffer, resuspended in PBS containing 1% BSA and 2 mM EDTA, and subjected to flow cytometry analysis by FACS Calibur (BD company). Data were analyzed using FlowJo software (version 7.6).

### Antibody production

Total RNA was extracted from 14A and16A murine hybridomas by QIAGEN RNeasy Mini reagent (QIAGEN). cDNA was synthesized by by SMARTer RACE reagent (CLONTECH). The primer used for reverse transcription was Oligo-dT. cDNA was used as PCR template to clone VH gene and VL gene. Universal primer A mix (CLONTECH) and 5’-GGGRCCARKG GATAGACHGATGG-3’ (designed according to the C segment of mouse IgG antibody heavy chain sequence) were used as cloning primers of VH gene. Universal primer A mix (CLONTECH) and 5’-5’
s-CTTCAGAGGA AGGGTGGAAACAGG-3’ (designed according to the C segment of mouse IgG antibody light chain sequence) were used as cloning primers of VL gene. VH and VL PCR fragments were sequenced by 3130XL ABI DNA sequencer.

Recombinant 14A and 16A antibodies were generated by jointing VH (fig. S3) to C region of mouse IgG1, and VH region to C region of mouse lambda chain. Full lengths of L and H genes were built into pcDNA3.1 expression vector (Invitrogen), as pcDNA3.1-L and pcDNA3.1-H, respectively. Pairs of pcDNA3.1-L and pcDNA3.1-H plasmids were transfected simultaneously by electroporation (Maxcyte). HEK293 cells were cultured in serum-free med media. Five days after electroporation, culture supernatant was combined, and antibody was purified by Protein A affinity chromatography column (GE Healthcare).

### Antigen and Purification

Peptides and glycopeptides were synthesized as described^11^ on an automated peptide synthesizer using fluorenylmethyloxycarbonyl (Fmoc)-protected amino acids as the building blocks, 6-chloro-benzotriazole-1-yl-oxy-tris-pyrrolidino-phosphonium hexafluorophosphate (PyClock) as the coupling reagent, and 2-chlorotrityl resin preloaded with glycine, loading capacity 0.59 mmol/g, as the solid support. In each coupling cycle PyClock and the Fmoc amino acid were used in 2.5-fold excess, and N-methylmorpholine (NMM) in 4.25-fold excess in N,N-dimethylformamide (DMF). Removal of the Fmoc group after each coupling was performed with 20% piperidine in DMF. For the glycopeptide preparation, the glycosylamino acid Fmoc-Thr(Ac_3_GalNAc)-OH was coupled in only 1.2-fold excess with the coupling reagent O-(7-Azabenzotriazol-1-yl)-N,N,N’,N’-tetramethyluronium hexafluorophosphate (HATU) and NMM. The peptide and glycopeptide were released from the resin with simultaneous deprotection by treatment with cocktail R (TFA/thioanisole/EDT/anisole, 90:5:3:2) and precipitated by cold ether.

The crude peptides were purified by preparative reversed-phase HPLC, followed by deacetylation with hydrazine. The mixture was shaken at room temperature for 30 min. The deacetylation reaction was monitored by analytical HPLC. The residue was neutralized with AcOH until pH 4, and subject to the preparative HPLC purification to give the de-acetylated product.

### Microarray analysis of mAbs

73 glycopeptides containing different regions of MUC1 tandem repeats (table S1) were used for microarray as described. 16A and 14A mAbs (mouse IgG1) were tested at different concentrations. VVA (Vicia Villosa Lectin) was used as control for detecting T_N_ antigen. The microarray analysis was performed as described (*28*).

### Production and purification of Fab fragments

The Fab fragments of anti-Muc1 mAbs 14A and 16A were expressed by Bac-to-Bac baculovirus expression system (Invitrogen). The heavy-chain (Hc; Fd fragment) gene of Fab includes the sequence of the variable domain of heavy chain, the human IgG1 CH1 and hinge, while the light-chain (Lc) gene of the Fab fragment includes the sequence of the variable domain of light chain and the human kappa constant domain. The Hc gene with a N-terminal gp67 signal peptide and Lc gene with a N-terminal honeybee melittin signal peptide were subcloned into pFastBac Dual vector (Invitrogen) containing two multiple cloning sites, with a Tev cleavage site followed by a 10×His tag at the C-terminus of the Hc to facilitate purification. The secreted Fab was captured by IMAC with a Ni^2+^-chelating affinity chromatography. The target protein was eluted with a buffer containing 20 mM Tris-HCl (pH 8.0), 150 mM NaCl, and 100 mM imidazole. The fractions were pooled and cleaved with Tev protease at 4 °C for 12 h to remove the C-terminal 10×His tag. The digested sample was loaded onto a Ni^2+^-chelating affinity chromatography column again to remove undigested Fab, and the cleaved product in the flow through was subjected to kappa affinity column (GE Healthcare). The eluted product from kappa affinity column was pooled and further purified by size-exclusion chromatography using a Superdex 200 10/300 GL column (GE Healthcare) equilibrated with a buffer containing 20 mM Tris-HCl (pH 8.0) and 50 mM NaCl.

Mutants of 16A (W34A, W34F, R32A and R32G) and 14A (G32R) were constructed using PCR with wide-type 16A or 14A expression vector as the template, respectively. The mutated sequences were confirmed by DNA sequencing. The mutants were expressed and purified using the same protocol as for wild-type Fab fragments.

### Protein Crystallization and Structure Determination

The purified Fab was concentrated to 10 mg/mL in 20 mM Tris-HCl (pH 8.0) and 50 mM NaCl buffer for crystallization. For the (glyco)peptide complex, antibody was mixed with (glyco)peptide in a 1:5 molar ratio overnight at 4 °C. The Fab alone and its ligand complex were screened respectively for crystallization using the hanging drop vapor diffusion at 20 °C with drops of 100 nL protein mixed with 100 nL well solution using a TTP LabTech Mosquito robot. Crystals were grown at 293K and typically appeared within 3 days. Optimal crystals of 16A apo were obtained in 0.1 M Tris-HCl pH 8.5, 1.0 M Lithium chloride, 20% PEG 6000. Interestingly, similar approximate conditions were found to generate the crystals of the 16A in complex with the glycopeptide Glyco-T, Glyco-S and Glyco-ST. Optimal crystals of 16A in complex with peptide were obtained in 0.2M di-sodium hydrogen phosphate, 20% PEG 3350. Optimal crystals of 14A apo were grown in 0.1 M Sodium cacodylate trihydrate pH 6.8, 0.2 M Sodium chloride, 25% PEG 6000. Crystals of 14A in complex with glycopeptide Glyco-T, Glyco-S and Glyco-ST were grown in 0.1 M Sodium citrate pH 5.5, 10% Isopropanol (v/v), 20% PEG 4000. The 14A in complex with peptide did not yield crystals in the screened conditions. Crystals were cryoprotected by a brief immersion in 70% well buffer and 30% glycerol, followed by immediate flash-cooling in liquid nitrogen.

X-ray diffraction data sets were collected on the BL19U beamline at the Shanghai Synchrotron Research Facility. All diffraction data were indexed, integrated, and scaled using HKL-2000. The structure was determined by the molecular replacement method with the program Phaser by using the variable and constant domains of the monoclonal antibody SYA/J6 (Protein Data Bank accession code 1M71) as an initial model. Model building was carried out using Coot, and refinement was implemented with the PHENIX program (*29*). The peptide was manually built into the difference density Fo-Fc map, followed by additional rounds of refinement of the complex in phenix.refine and manual building cycles in Coot. The final models were validated using MolProbity. Data collection and refinement statistics are provided in table S2. All structural visualizations were generated with PyMOL. For the Fab, the residues were renumbered according to the Kabat scheme.

### Surface Plasmon Resonance (SPR) Measurement of Ab Binding Affinity

Interactions of RPAPGSTAPPAHG, RPAPGS(GalNAc)TAPPAHG, RPAPGST(GalNAc)APPAHG and RPAPGS(GalNAc)T(GalNAc)APPAHG with immobilized antibody 14A and 16A were determined by surface plasmon resonance on a Biacore T200 (GE Healthcare) instrument. Antibody 14A and 16A were immobilized on a CM5 chip until reaching 4000 response units. A reference channel was immobilized with ethanolamine, respectively. Immobilizations were carried out at protein concentrations of 25 μg/ml in 10 mM acetate pH 5.0 by using an amine coupling kit supplied by the manufacturer. Measurements were carried out at 25°C in 10 mM HEPES, pH 7.4 containing 150 mM NaCl and 0.005% surfactant P20 at a flow rate of o 30 μl/min. The association time was 180 s and dissociation time was 300 s. The surface was regenerated by 1 mM NaOH solution. Data were analyzed with BIA evaluation software (GE Healthcare).

### Molecular dynamics simulation and analysis

Based on the structures of the antibody in complex with the MUC1 peptides, we generated the initial model for MD simulations. Considering the simulation efficiency, only the Fv domain of each antibody that are in close contact with the MUC1 peptides was kept in the simulations. The peptide [CH3-(C=O)-]PGSTAPPAHG[-(C=O)-NH2] was used in simulations. The protonation states of all the residues at pH 7.0 were predicted with PROPKA (*30*) module in Schrodinger Suite v2019-1 (http://www.schrodinger.com). In order to investigate the glycosylation effects, six systems were generated by varying the presence of glycosyl groups that can be attached to the Ser/Thr (see table S3). Each structure then was solvated into a truncated dodecahedron box by adding water molecules and 0.15 M NaCl. The buffering distance of proteins to the box boundary is at least 12 Å.

All simulations were performed using Gromacs 2018.4 package (*31,32*) with Amber ff14SB force field (*33*) and TIP3P explicit water model. The CLYCAM_06j-1 parameters (*34*) and non-uniform 1-4 scaling factor strategy were employed. Prior to long MD simulations, unrestrained geometry optimization was carried out, followed by a two-stage equilibration. The systems were equilibrated with constant volume at 300 K (NVT) in stage one, and then under constant pressure of 1 atm at 300 K in stage two. The v-rescale temperature coupling scheme (*35*) and Parrinello-Rahman pressure coupling scheme (*36*) were used to control the temperature and pressure during simulations. The long range electrostatic interaction was calculated with particle-mesh-Ewald (PME) method (*37*), and the cutoff of short-range electrostatic interaction is 12 Å. After equilibration, production simulations were performed for 300 ns in NPT ensemble with the condition of 300 K and 1 atm pressure. The time step was 2 fs, and the system coordinates were saved every 0.1 ns. The trajectories of last 250 ns simulations were used for dynamics analysis.

The structural deviation and fluctuations (i.e., the Root-Mean-Squared-Deviation or RMSD; Root-Mean-Squared-Fluctuation, RMSF) were calculated with programs included in Gromacs package. The structural features, such as distances, angles and dihedral angles were measured using VMD scripts (*38*). The hydrogen bonds were analyzed with the HBonds Plugin in VMD. The hydrogen bond Donor-Acceptor distance cutoff was set to 3.5 Å, the angle cutoff was 30 degrees.

### Statistical analysis

Not applicable.

## Acknowledgments

We thank IA Wilson for advice, the staffs of beamlines BL19U at the National Center for Protein Sciences Shanghai and Shanghai Synchrotron Radiation Facility for assistance in data collection. We also thank the Discovery Technology Platform of Shanghai Institute for Advanced Immunochemical Studies (SIAIS), ShanghaiTech University, for equipment service.

## Author contributions

Conceptualization: D.Z., L.X., and H.L.

Methodology: Y.H., J.N., D.P., C.F., K.S., B.M., U.W., and Y.Z.

Investigation: Y.H., J.N., D.P., C.F., K.S.

Visualization: Y.H., J.N., D.P., C.F., K.S., U.W.

Supervision: D.Z., L.X.

Writing—original draft: D.Z., Y.H. and H.L.

Writing—review & editing: D.Z., Y.H. and H.L.

## Data and materials availability

The crystal structural data has been deposited with PDB ID as follows: 16A, 7V3Q; 16A/Peptide, 7V4W; 16A/GlycoT, 7V64; 16A/GlycoST, 7V7K; 14A/GlycoS, 7VAZ; 14A/GlycoT, 7V8Q; and 14A/GlycoST, 7VAC.

## References

1. A. P. Corfield, Mucins: A biologically relevant glycan barrier in mucosal protection. Biochim. Biophys. Acta. Gen. Subj. 1850, 236–252 (2015).

2. K. O. Saunders, N. I. Nicely, K. Wiehe, M. Bonsignori, R. R. Meyerhoff, R. Parks, W. E. Walkowicz, B. Aussedat, N. R. Wu, F. Cai, Y. Vohra, P. K. Park, A. Eaton, E. P. Go, L. L. Sutherland, R. M. Scearce, D. H. Barouch, R. Zhang, T. Von Holle, R. G. Overman, K. Anasti, R. W. Sanders, M. A. Moody, T. B. Kepler, B. Korber, H. Desaire, S. Santra, N. L. Letvin, G. J. Nabel, D. C. Montefiori, G. D. Tomaras, H.-X. Liao, S. M. Alam, S. J. Danishefsky, B. F. Haynes, Vaccine Elicitation of High Mannose-Dependent Neutralizing Antibodies against the V3-Glycan Broadly Neutralizing Epitope in Nonhuman Primates. Cell Rep. 18, 2175–2188 (2017).

3. T. Freund Natalia, H. Wang, L. Scharf, L. Nogueira, A. Horwitz Joshua, Y. Bar-On, J. Golijanin, A. Sievers Stuart, D. Sok, H. Cai, C. Cesar Lorenzi Julio, A. Halper-Stromberg, I. Toth, A. Piechocka-Trocha, B. Gristick Harry, J. van Gils Marit, W. Sanders Rogier, L.-X. Wang, S. Seaman Michael, R. Burton Dennis, A. Gazumyan, D. Walker Bruce, P. West Anthony, J. Bjorkman Pamela, C. Nussenzweig Michel, Coexistence of potent HIV-1 broadly neutralizing antibodies and antibody-sensitive viruses in a viremic controller. Sci. Transl. Med. 9, eaal2144 (2017).

4. Y. Watanabe, D. Allen Joel, D. Wrapp, S. McLellan Jason, M. Crispin, Site-specific glycan analysis of the SARS-CoV-2 spike. Science 369, 330–333 (2020).

5. D. Zhou, X. Tian, R. Qi, C. Peng, W. Zhang, Identification of 22 N-glycosites on spike glycoprotein of SARS-CoV-2 and accessible surface glycopeptide motifs: Implications for vaccination and antibody therapeutics. Glycobiology 31, 69–80 (2020).

6. V. Lakshminarayanan, P. Thompson, M. A. Wolfert, T. Buskas, J. M. Bradley, L. B. Pathangey, C. S. Madsen, P. A. Cohen, S. J. Gendler, G.-J. Boons, Immune recognition of tumor-associated mucin MUC1 is achieved by a fully synthetic aberrantly glycosylated MUC1 tripartite vaccine. Proc. Natl. Acad. Sci. U S A. 109, 261–266 (2012).

7. H. Cai, R.-S. Zhang, J. Orwenyo, J. Giddens, Q. Yang, C. C. LaBranche, D. C. Montefiori, L.- Wang, Synthetic HIV V3 Glycopeptide Immunogen Carrying a N334 N-Glycan Induces Glycan-Dependent Antibodies with Promiscuous Site Recognition. J. Med. Chem. 61, 10116–10125 (2018).

8. D. Zhou, L. Xu, W. Huang, T. Tonn, Epitopes of MUC1 Tandem Repeats in Cancer as Revealed by Antibody Crystallography: Toward Glycopeptide Signature-Guided Therapy. Molecules 23, (2018).

9. M. A. Cheever, J. P. Allison, A. S. Ferris, O. J. Finn, B. M. Hastings, T. T. Hecht, I. Mellman, S. A. Prindiville, J. L. Viner, L. M. Weiner, L. M. Matrisian, The Prioritization of Cancer Antigens: A National Cancer Institute Pilot Project for the Acceleration of Translational Research. Clin. Cancer Res. 15, 5323–5337 (2009).

10. M. P. Torres, S. Chakraborty, J. Souchek, S. K. Batra, Mucin-based Targeted Pancreatic Cancer Therapy. Curr. Pharm. Des.18, 2472–2481 (2012).

11. S. Cascio, O. J. Finn, Intra- and Extra-Cellular Events Related to Altered Glycosylation of MUC1 Promote Chronic Inflammation, Tumor Progression, Invasion, and Metastasis. Biomolecules 6, 39 (2016).

12. E. P. Bennett, U. Mandel, H. Clausen, T. A. Gerken, T. A. Fritz, L. A. Tabak, Control of mucin-type O-glycosylation: A classification of the polypeptide GalNAc-transferase gene family. Glycobiology 22, 736–756 (2011).

13. T. Ju, Y. Wang, R. P. Aryal, S. D. Lehoux, X. Ding, M. R. Kudelka, C. Cutler, J. Zeng, J. Wang, X. Sun, J. Heimburg-Molinaro, D. F. Smith, R. D. Cummings, Tn and sialyl-Tn antigens, aberrant O-glycomics as human disease markers. Proteomics Clin. Appl. 7, 618–631 (2013).

14. A. Schietinger, M. Philip, B. A. Yoshida, P. Azadi, H. Liu, S. C. Meredith, H. Schreiber, A Mutant Chaperone Converts a Wild-Type Protein into a Tumor-Specific Antigen. Science 314, 304–308 (2006).

15. J. Zeng, R. P. Aryal, K. Stavenhagen, C. Luo, R. Liu, X. Wang, J. Chen, H. Li, Y. Matsumoto Wang, J. Wang, T. Ju, R. D. Cummings, Cosmc deficiency causes spontaneous autoimmunity by breaking B cell tolerance. Sci. Adv. 7, eabg9118 (2021).

16. P. Braun, G. M. Davies, M. R. Price, P. M. Williams, S. J. B. Tendler, H. Kunz, Effects of glycosylation on fragments of tumour associated human epithelial mucin MUC1. Bioorg. Med. Chem. 6, 1531–1545 (1998).

17. S. Dziadek, C. Griesinger, H. Kunz, U. M. Reinscheid, Synthesis and Structural Model of an α(2,6)-Sialyl-T Glycosylated MUC1 Eicosapeptide under Physiological Conditions. Chemistry – A European Journal 12, 4981–4993 (2006).

18. N. Martínez-Sáez, J. Castro-López, J. Valero-González, D. Madariaga, I. Compañón, V. J. Somovilla, M. Salvadó, J. L. Asensio, J. Jiménez-Barbero, A. Avenoza, J. H. Busto, G. J. L. Bernardes, J. M. Peregrina, R. Hurtado-Guerrero, F. Corzana, Deciphering the Non-Equivalence of Serine and Threonine O-Glycosylation Points: Implications for Molecular Recognition of the Tn Antigen by an anti-MUC1 Antibody. Angew. Chem. Int. Ed. Engl. 54, 9830–9834 (2015).

19. M. Movahedin, T. M. Brooks, N. T. Supekar, N. Gokanapudi, G.-J. Boons, C. L. Brooks, Glycosylation of MUC1 influences the binding of a therapeutic antibody by altering the conformational equilibrium of the antigen. Glycobiology 27, 677–687 (2017).

20. H. Wakui, Y. Tanaka, T. Ose, I. Matsumoto, K. Kato, Y. Min, T. Tachibana, M. Sato, K. Naruchi, F. G. Martin, H. Hinou, S. I. Nishimura, A straightforward approach to antibodies recognising cancer specific glycopeptidic neoepitopes. Chem. Sci. 11, 4999–5006 (2020).

21. W. Song, E. S. Delyria, J. Chen, W. Huang, J. S. Lee, E. A. Mittendorf, N. Ibrahim, L. G. Radvanyi, Y. Li, H. Lu, H. Xu, Y. Shi, L.-X. Wang, J. A. Ross, S. P. Rodrigues, I. C. Almeida, X. Yang, J. Qu, N. S. Schocker, K. Michael, D. Zhou, MUC1 glycopeptide epitopes predicted by computational glycomics. Int. J. Oncol. 41, 1977–1984 (2012).

22. C. L. Brooks, A. Schietinger, S. N. Borisova, P. Kufer, M. Okon, T. Hirama, C. R. MacKenzie, L.-X. Wang, H. Schreiber, S. V. Evans, Antibody recognition of a unique tumor-specific glycopeptide antigen. Proc. Natl. Acad. Sci. U S A. 107, 10056–10061 (2010).

23. P. Sharma, V. V. V. R. Marada, Q. Cai, M. Kizerwetter, Y. He, S. P. Wolf, K. Schreiber, H. Clausen, H. Schreiber, D. M. Kranz, Structure-guided engineering of the affinity and specificity of CARs against Tn-glycopeptides. Proc. Natl. Acad. Sci. U S A. 117, 15148–15159 (2020).

24. K. R. Trabbic, K. A. Kleski, M. Shi, J.-P. Bourgault, J. M. Prendergast, D. T. Dransfield, P. R. Andreana, Production of a mouse monoclonal IgM antibody that targets the carbohydrate Thomsen-nouveau cancer antigen resulting in in vivo and in vitro tumor killing. Cancer Immunol. Immunother. 67, 1437–1447 (2018).

25. A. Babino, O. Pritsch, P. Oppezzo, R. Du Pasquier, A. Roseto, E. Osinaga, P. M. Alzari, Molecular Cloning of a Monoclonal Anti-Tumor Antibody Specific for the Tn Antigen and Expression of an Active Single-Chain Fv Fragment. Hybridoma 16, 317–324 (1997).

26. R. L. Brady, D. J. Edwards, R. E. Hubbard, J. S. Jiang, G. Lange, S. M. Roberts, R. J. Todd, J. R. Adair, J. S. Emtage, D. J. King, D. C. Low, Crystal structure of a chimeric Fab′ fragment of an antibody binding tumour cells. J. Mol. Biol.227, 253–264 (1992).

27. R. Henderson, B. E. Watts, H. N. Ergin, K. Anasti, R. Parks, S.-M. Xia, A. Trama, H.-X. Liao, K. O. Saunders, M. Bonsignori, K. Wiehe, B. F. Haynes, S. M. Alam, Selection of immunoglobulin elbow region mutations impacts interdomain conformational flexibility in HIV-1 broadly neutralizing antibodies. Nat. Commun. 10, 654 (2019).

28. C. Pett, H. Cai, J. Liu, B. Palitzsch, M. Schorlemer, S. Hartmann, N. Stergiou, M. Lu, H. Kunz, E. Schmitt, U. Westerlind, Microarray Analysis of Antibodies Induced with Synthetic Antitumor Vaccines: Specificity against Diverse Mucin Core Structures. Chemistry – A European Journal 23, 3875–3884 (2017).

29. D. Liebschner, P. V. Afonine, M. L. Baker, G. Bunkóczi, V. B. Chen, T. I. Croll, B. Hintze, L.-W. Hung, S. Jain, A. J. McCoy, N. W. Moriarty, R. D. Oeffner, B. K. Poon, M. G. Prisant, R. J. Read, J. S. Richardson, D. C. Richardson, M. D. Sammito, O. V. Sobolev, D. H. Stockwell, T. C. Terwilliger, A. G. Urzhumtsev, L. L. Videau, C. J. Williams, P. D. Adams, Macromolecular structure determination using X-rays, neutrons and electrons: recent developments in Phenix. Acta Crystallographica Section D-Structural Biology 75, 861–877 (2019).

30. M. H. M. Olsson, C. R. Søndergaard, M. Rostkowski, J. H. Jensen, PROPKA3: Consistent Treatment of Internal and Surface Residues in Empirical pKa Predictions. J. Chem. Theory. Comput. 7, 525–537 (2011).

31. S. Pronk, S. Páll, R. Schulz, P. Larsson, P. Bjelkmar, R. Apostolov, M. R. Shirts, J. C. Smith, AP. M. Kasson, D. van der Spoel, B. Hess, E. Lindahl, GROMACS 4.5: a high-throughput and highly parallel open source molecular simulation toolkit. Bioinformatics 29, 845–854 (2013).

32. M. J. Abraham, T. Murtola, R. Schulz, S. Páll, J. C. Smith, B. Hess, E. Lindahl, GROMACS: High performance molecular simulations through multi-level parallelism from laptops to supercomputers. SoftwareX 1-2, 19–25 (2015).

33. J. A. Maier, C. Martinez, K. Kasavajhala, L. Wickstrom, K. E. Hauser, C. Simmerling, ff14SB: Improving the Accuracy of Protein Side Chain and Backbone Parameters from ff99SB. J. Chem. Theory. Comput. 11, 3696–3713 (2015).

34. K. N. Kirschner, A. B. Yongye, S. M. Tschampel, J. González-Outeiriño, C. R. Daniels, B. L. Foley, R. J. Woods, GLYCAM06: A generalizable biomolecular force field. Carbohydrates. J. Comput. Chem. 29, 622–655 (2008).

35. G. Bussi, D. Donadio, M. Parrinello, Canonical sampling through velocity rescaling. J. Chem. Phys. 126, 014101 (2007).

36. M. Parrinello, A. Rahman, Polymorphic transitions in single crystals: A new molecular dynamics method. Journal of Applied Physics 52, 7182–7190 (1981).

37. T. Darden, D. York, L. Pedersen, Particle mesh Ewald: An N·log(N) method for Ewald sums in large systems. J. Chem. Phys. 98, 10089–10092 (1993).

38. W. Humphrey, A. Dalke, K. Schulten, VMD: Visual molecular dynamics. Journal of Molecular Graphics 14, 33–38 (1996).

