## Supplemental online materials for "Structural basis of a high-affinity antibody binding to glycoprotein region with consecutive glycosylation sites"

**This PDF file includes:**

Figs. S1 to S3

Tables S1 to S3

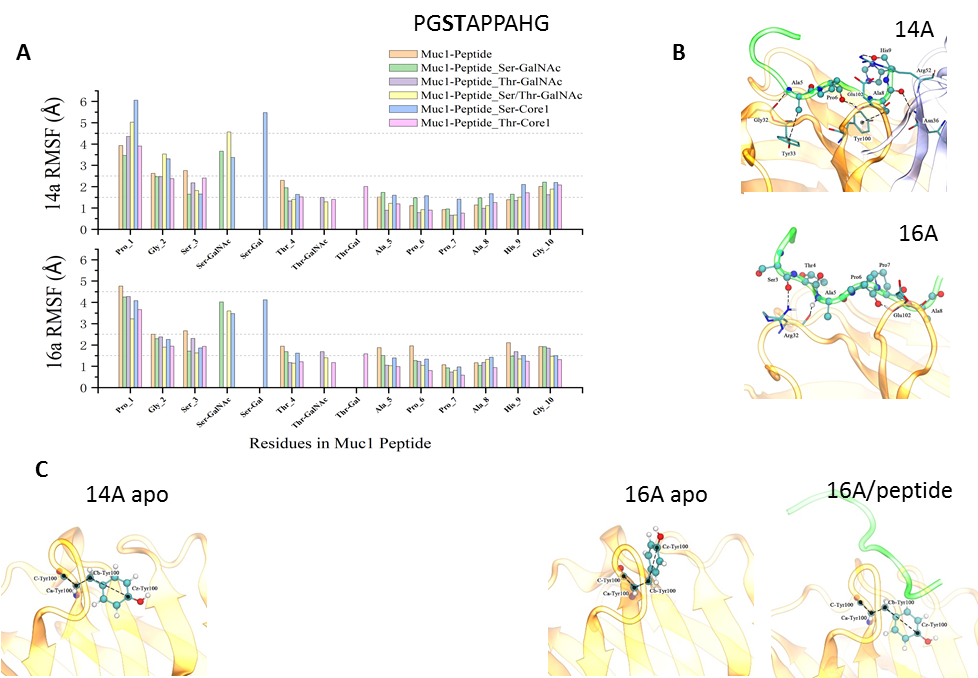

**Fig. S1. Dynamics and interactions revealed from molecular dynamics simulations.** (**A**) Peptide residue flucutations for peptides with different glycosylations in complex with antibodies 14A (upper panel) and 16A (lower panel). (**B**) The interactions between MUC1-peptides and antibodies. The hydrogen bonds formed between Arg32 in heavy chain of 16A and Ser3 in peptide is not observed for 14A/peptide system. The peptide residues is presented with CPK, and antibody was in licorice representation. (**C**) The conformations of Tyr100 in heavy chain of 14A and 16A, in apo and peptide bound structures. The co-crystallization of 14A and non-glycosylated peptide was not achieved experimentally.

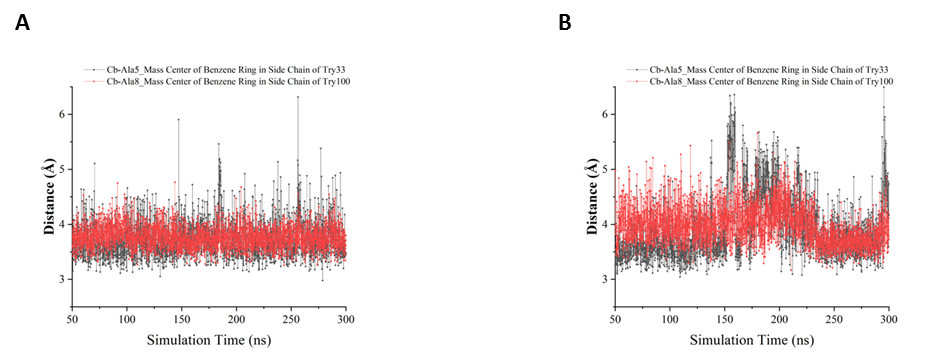

**Fig. S2.** **The distances between peptide residues and the antibodies.** (**A**) the distance between Cβ of Ala5 and mass center of benzene ring of Tyr33 and the distance between Cβ of Ala8 and mass center of benzene ring of Tyr100 in 14A/peptide. (**B**) same distance plots for 16A/peptide system.

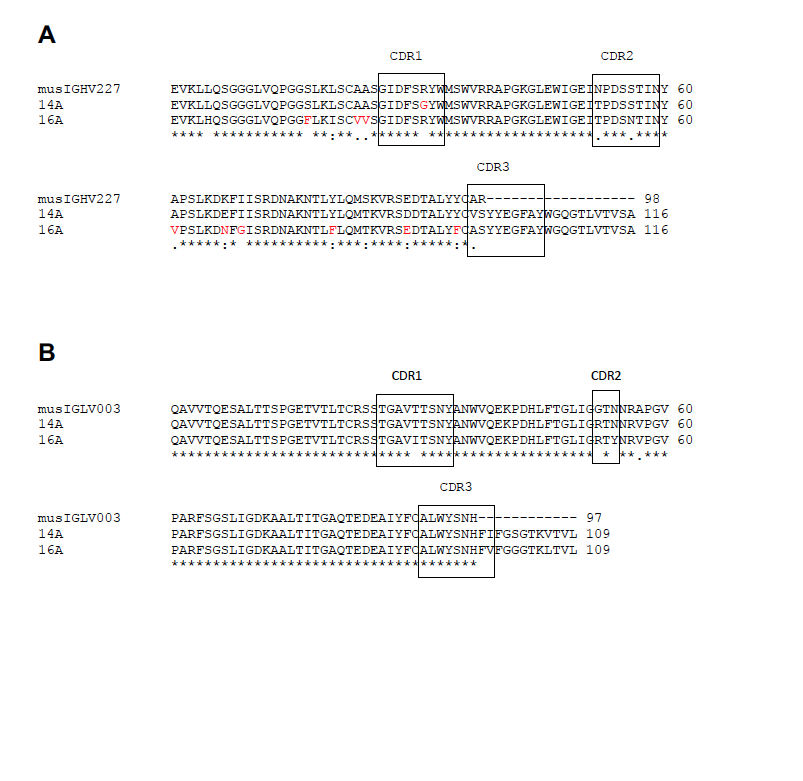

**Fig. S3.** Sequence alignment of VL and VH genes of 14A and 16A with germline IGHV (**A**) and IgLV (**B**) genes.

Table S1. Glycopeptides used in microarray experiments.

Glycosylated amino acids are labelled as red color. Black color represent amino acid sequences. * indicates the site of glycosylation. Translucent amino acid sequences are not present in the synthesized glycopeptides.

T_N_, GalNAc

T, Galβ1,3GalNAc

ST, NeuAcα2,3Galβ1,3GalNAc

ST_N_, NeuAcα2,6GalNAc

T2C3, Galβ1,4GlcNAcβ1,3GalNAc

βGlcNAc

| 1 | PAHGVT*(T_N_)SAPDTRPAPGSTA |
| --- | --- |
| 2 | PAHGVTSAPDT*(T_N_)RPAPGSTA |
| 3 | PAHGVTSAPDTRPAPGST*(T_N_)A |
| 4 | PAHGVT*(T_N_)SAPDT*(T_N_)RPAPGSTA |
| 5 | PAHGVT*(T_N_)SAPDTRPAPGST*(T_N_)A |
| 6 | PAHGVTSAPDT*(T_N_)RPAPGST*(T_N_)A |
| 7 | PAHGVT*(T_N_)SAPDT*(T_N_)RPAPGST*(T_N_)A |
| 8 | PAHGVT*(T)SAPDTRPAPGSTA |
| 9 | PAHGVTSAPDT*(T)RPAPGSTA |
| 10 | PAHGVTSAPDTRPAPGST*(T)A |
| 11 | PAHGVT*(T)SAPDT*(T)RPAPGSTA |
| 12 | PAHGVT*(T)SAPDTRPAPGST*(T)A |
| 13 | PAHGVTSAPDT*(T)RPAPGST*(T)A |
| 14 | PAHGVT*(T)SAPDT*(T)RPAPGST*(T)A |
| 15 | PGSTAPPAHGVTSAPDT*(T)RPA |
| 16 | APDT*(T)RPAPGSTAPPAǀHGVTSA |
| 17 | APDT*(T)RPA |
| 18 | PAHGVT*(T2C3)SA |
| 19 | PAHGVT*(ST)SAPDTRPAPGSTA |
| 20 | PAHGVTSAPDT*(ST)RPAPGSTA |
| 21 | PAHGVTSAPDTRPAPGST*(ST)A |
| 22 | PAHGVT*(ST)SAPDT*(ST)RPAPGSTA |
| 23 | PAHGVT*(ST)SAPDTRPAPGST*(ST)A |
| 24 | PAHGVTSAPDT*(ST)RPAPGST*(ST)A |
| 25 | PAHGVT*(ST)SAPDT*(ST)RPAPGST*(ST)A |
| 26 | PAHGVTSAPDTRPAPGSTAP |
| 27 | PAHGVT*(ST_N_)SAPDTRPAPGSTAP |
| 28 | PAHGVTSAPDT*(ST_N_)RPAPGSTAP |
| 29 | PAHGVTSAPDTRPAPGST*(ST_N_)AP |
| 30 | PAHGVT*(ST_N_)SAPDTRPAPGST*(ST_N_)AP |
| 31 | PAHGVT*(ST_N_)SAPDTRPAP |
| 32 | GSTAPPAǀHGVT*(ST_N_)SAP |
| 33 | PAHGVT*( ST_N_)SAPDTRPAPGST*(T_N_)AP |
| 34 | PAHGVT*( ST_N_)SAPDT*(T_N_)RPAPGST*(T_N_)AP |
| 35 | PAHGVT*( ST_N_)SAPDTRPAPGST*(T_N_)APPAHGVT*(ST_N_)SAPDTRPAPGST*(T_N_)AP |
| 36 | PAHGVT*(T_N_)SAPDTRPAPGSTAP |
| 37 | PAHGVTS*(T_N_)APDTRPAPGSTAP |
| 38 | PAHGVTSAPDTRPAPGS*(T_N_)TAP |
| 39 | PAHGVTSAPDT*(T_N_)RPAPGST*(T_N_)AP |
| 40 | APDTRPAPGS*(T_N_)TAP |
| 41 | PAHGVT*(T)SAPDTRPAPGSTAP |
| 42 | PAHGVTS*(T)APDTRPAPGSTAP |
| 43 | PAHGVTSAPDT*(T)RPAPGSTAP |
| 44 | PAHGVTSAPDTRPAPGS*(T)TAP |
| 45 | PAHGVTSAPDTRPAPGST*(T)AP |
| 46 | PAHGVT*(T)SAPDTRPAP |
| 47 | PAHGVTS*(T)APDTRPAP |
| 48 | HGVTSAPDTRPAPGSTAPPA |
| 49 | HGVTSAPDT*(T_N_)RPAPGSTAPPA |
| 50 | HGVTSAPDT*(T_N_)RPAPGS*(T_N_)TAPPA |
| 51 | HGVTSAPDT*(T_N_)RPAPGST*(T_N_)APPA |
| 52 | HGVTSAPDT*(T_N_)RPAPGS*(T_N_)T*(T_N_)APPA |
| 53 | HGVTSAPDT*(T)RPAPGS*(T)T*(T)APPA |
| 54 | HGVTSAPDT*(bGlcNAc)RPAPGS*(T_N_)T*(T_N_)APPA |
| 55 | HGVTSAPDTRPAPGS*(T_N_)T*(T_N_)APPA |
| 56 | HGVTSAPDTRPAPGS*(T)T*(T)APPA |
| 57 | HGVT*(T_N_)S*(T_N_)APDT*(T_N_)RPAPGS*(T_N_)T*(T_N_)APPA |
| 58 | HGVTSAPDTRPAPGS*(T_N_)TAPPA |
| 59 | HGVTSAPDTRPAPGST*(T_N_)APPA |
| 60 | HGVTSA |
| 61 | HGVTSAPDTRPA |
| 62 | HGVTSAPDT*(T_N_)RPA |
| 63 | HGVT*(T_N_)S*(T_N_)APDT*(T_N_)RPA |
| 64 | APDTRPAPGSTAPPA |
| 65 | APDT*(T_N_)RPAPGSTAPPA |
| 66 | APDT*(βGlcNAc)RPAPGSTAPPA |
| 67 | APDTRPAPGS*(T_N_)T*(T_N_)APPA |
| 68 | APDT*(T_N_)RPAPGS*(T_N_)T*(T_N_)APPA |
| 69 | APDTRPA |
| 70 | APDT*(T_N_)RPA |
| 71 | APGSTAPPA |
| 72 | APGS*(T_N_)TAPPA |
| 73 | APGS*(T_N_)T*(T_N_)APPA |

Table S2. Data collection and refinement statistics

|  | 16A | 16A/Peptide | 16A/GlycoT | 16A/GlycoST | 14A/GlycoS | 14A/GlycoT | 14A/GlycoST |
| --- | --- | --- | --- | --- | --- | --- | --- |
| **Data collection** |  | | | |  |  |  |
| Space group | P2_1_2_1_2_1_ | P12_1_1 | P2_1_2_1_2_1_ | P2_1_2_1_2_1_ | P12_1_1 | P2_1_2_1_2_1_ | P2_1_2_1_2_1_ |
| Cell dimensions |  | | | |  |  |  |
| a, b, c (Å) | 43.6, 126.7, 158.2 | 39.6, 63.4, 88.7 | 39.5, 81.2, 137.3 | 39.1, 136.6, 80.9 | 46.6, 226.6, 71.1 | 42.9, 207.4, 224.7 | 42.9, 206.8, 225.4 |
| α, β, γ (Å) | 90, 90, 90 | 90, 97.1, 90 | 90, 90, 90 | 90, 90, 90 | 90, 107.7, 90 | 90, 90, 90 | 90, 90, 90 |
| Resolution (Å)^*^ | 48.68-2.98  (3.09-2.98) | 39.25-2.10  (2.18-2.10) | 40.61-1.56  (1.62-1.56) | 39.70-2.20  (2.28-2.20) | 44.36-2.73  (2.83-2.73) | 39.63-3.20  (3.32-3.20) | 40.69-3.50  (3.63-3.50) |
| No. of observed reflections | 308,245 | 262,777 | 1,759,168 | 151,736 | 960,221 | 2,535,056 | 1,874,543 |
| Completeness(%) | 97.7 (92.3) | 91.5 (54.4) | 98.1 (88.9) | 93.4(77.5) | 98.4(84.9) | 98.4(96.2) | 99.7(99.5) |
| R*_meas_*(%)^a^ | 18.4(72.8) | 12.9(39.0) | 10.4(85.5) | 12.9(73.1) | 13.1(48.5) | 31.5(75.4) | 16.4(84.1) |
| R*_pim_*(%)^b^ | 8.6(35.0) | 6.8(24.1) | 2.9(25.8) | 7.2(49.0) | 5.0(20.8) | 9.1(24.9) | 4.8(25.3) |
| <I/σ(I)> | 9.1(2.1) | 25.3(3.4) | 23.7(2.2) | 10.5(1.2) | 10.8(2.0) | 5.0(2.0) | 11.7(2.0) |
| CC_1/2_^c^ | 0.970(0.707) | 0.964(0.848) | 0.999(0.849) | 0.994(0.580) | 0.997(0.846) | 0.989(0.766) | 0.998(0.816) |
| Redundancy | 4.1(3.9) | 1.8(1.3) | 6.4(5.4) | 1.6(1.1) | 6.6(5.2) | 6.3(4.3) | 6.2(5.7) |
| Wilson B (Å^2^) | 42.9 | 29.8 | 15.5 | 27.2 | 41.8 | 72.2 | 62.2 |
| **Refinement statistics** |  | | | |  |  |  |
| Reflections used in refinement | 18,223 (1,686) | 23,401 (1,379) | 62,756 (5,598) | 21,331 (1,731) | 36,277 (3,078) | 33,724 (3,260) | 26,442 (2,588) |
| Non-hydrogen atoms | 6,556 | 3,574 | 3,835 | 3,605 | 9,884 | 10,028 | 10,013 |
| Macromolecules | 6,556 | 3,292 | 3,298 | 3,333 | 9,864 | 9,986 | 9,929 |
| Ligands | - | - | 14 | 28 | 14 | 42 | 84 |
| Solvent | - | 282 | 523 | 244 | 6 | - | - |
| Average B-factors(Å^2^) | 37.7 | 31.6 | 20.2 | 29.3 | 42.8 | 65.4 | 55.2 |
| Macromolecules | 37.7 | 31.3 | 18.1 | 29.1 | 42.8 | 65.4 | 55.1 |
| Ligands | - | - | 22.2 | 29.5 | 36.1 | 67.8 | 70.1 |
| Solvent | - | 35.3 | 33.2 | 32.4 | 22.3 | - | - |
| R*_work_*/R*_free_*(%)^d^ | 21.8/26.4 | 18.0/23.3 | 18.0/20.3 | 20.1/25.1 | 20.9/26.4 | 20.0/24.4 | 17.1/21.9 |
| Rmsd bond length(Å) | 0.005 | 0.005 | 0.007 | 0.005 | 0.008 | 0.004 | 0.003 |
| Rmsd bond angle(°) | 0.84 | 0.84 | 0.92 | 0.89 | 1.16 | 0.86 | 0.66 |
| Ramachandran plot |  | | | |  |  |  |
| Favored(%) | 95.3 | 97.0 | 97.9 | 97.0 | 96.5 | 96.0 | 96.6 |
| Allowed(%) | 4.7 | 3.0 | 2.1 | 3.0 | 3.5 | 4.0 | 3.4 |
| Outliers(%) | 0 | 0 | 0 | 0 | 0 | 0 | 0 |

*Statistics for the highest-resolution shell are shown in parentheses.

^a^$R_{meas}=\frac{{\sum_{hkl} [\frac{N\left( hkl \right)}{N\left( hkl \right)-1}]}^{\frac{1}{2}}\times\sum_{i} \left| I_{i}\left( hkl \right)-\left\langle I(hkl) \right\rangle\right|}{\sum_{hkl} \sum_{i} I_{i}(hkl)}$

^b^ $R_{pim}=\frac{\sum_{hkl} \left[ \frac{1}{N\left( hkl \right)-1} \right]^{\frac{1}{2}}\times\sum_{i} \left| I_{i}\left( hkl \right)-\left\langle I(hkl) \right\rangle\right|}{\sum_{hkl} \sum_{i} I_{i}(hkl)}$

^c^ ${CC}_{\frac{1}{2}}=\frac{\sum(x-\left\langle x \right\rangle)(y-\left\langle y \right\rangle)}{\left[ \sum{(x-\left\langle x \right\rangle)}^{2}\sum{(y-\left\langle y \right\rangle)}^{2} \right]^{\frac{1}{2}}}$

*^d^ R_work_* $=\frac{\sum\left| \left| F_{obs} \right|-\left| F_{calc} \right| \right|}{\sum\left| F_{obs} \right|}$; *R_free_* is defined as *R_work_* calculated from 5% of the reflections that were excluded from refinement.

Table S3. Hydrogen Bond Analysis of MUC1-Peptide/Antibodies (14A/16A) systems

|  | **MUC1-Peptide** | | | **MUC1-Peptide_Ser-GalNAc** | | | **MUC1-Peptide_Ser-Core1** | | | **MUC1-Peptide_Ser/Thr-GalNAc** | | | **MUC1-Peptide_Thr-GalNAc** | | | **MUC1-Peptide_Thr-Core1** | | |
| --- | --- | --- | --- | --- | --- | --- | --- | --- | --- | --- | --- | --- | --- | --- | --- | --- | --- | --- |
| **14A** | Donor | Acceptor | Occupancy | Donor | Acceptor | Occupancy | Donor | Acceptor | Occupancy | Donor | Acceptor | Occupancy | Donor | Acceptor | Occupancy | Donor | Acceptor | Occupancy |
|  | H_Glu102-Main | M_Pro6-Main | 80.14% | H_Glu102-Main | M_Pro6-Main | 82.97% | M_Ala5-Main | H_Gly32-Main | 77.30% | M_Ala5-Main | H_Gly32-Main | 83.73% | M_Ala5-Main | H_Gly32-Main | 84.53% | M_Ala5-Main | H_Gly32-Main | 83.49% |
|  | L_Asn36-Side | M_Ala8-Main | 71.18% | M_Ala5-Main | H_Gly32-Main | 81.33% | H_Glu102-Main | M_Pro6-Main | 77.14% | H_Glu102-Main | M_Pro6-Main | 81.18% | H_Glu102-Main | M_Pro6-Main | 79.74% | H_Glu102-Main | M_Pro6-Main | 74.66% |
|  | M_Ala5-Main | H_Gly32-Main | 69.74% | L_Asn36-Side | M_Ala8-Main | 58.71% | L_Asn36-Side | M_Ala8-Main | 57.87% | L_Asn36-Side | M_Ala8-Main | 51.48% | L_Asn36-Side | M_Ala8-Main | 54.36% | L_Asn36-Side | M_Ala8-Main | 58.63% |
|  | L_Arg52-Side | M_His9-Main | 30.98% | L_Arg52-Side | M_His9-Main | 30.46% | L_Arg52-Side | M_His9-Main | 32.69% | H_Trp34-Side | M_Thr-GalNAc-Side | 36.17% | H_Trp34-Side | M_Thr-GalNAc-Side | 35.85% | L_Arg52-Side | M_His9-Main | 40.21% |
|  | L_Arg52-Main | M_Ala8-Main | 17.07% | L_Arg52-Side | M_Pro7-Main | 15.71% | M_Ser-GalNAc-Side | H_Tyr101-Side | 13.39% | L_Arg52-Side | M_His9-Main | 30.70% | L_Arg52-Side | M_His9-Main | 28.70% | H_Trp34-Side | M_Thr-GalNAc-Side | 27.38% |
|  | L_Arg52-Side | M_Gly10-Main | 15.51% | M_Ser-GalNAc-Side | H_Asp29-Side | 14.19% | L_Arg52-Side | M_Pro7-Main | 10.31% | M_Ser-GalNAc-Side | H_Asp29-Side | 18.15% | L_Arg52-Side | M_Pro7-Main | 15.83% | M_Thr-GalNAc-Side | H_Asp55-Side | 18.31% |
|  |  |  |  | M_Ser-GalNAc-Side | H_Tyr101-Side | 11.67% |  |  |  | L_Arg52-Side | M_Pro7-Main | 16.27% |  |  |  | L_Arg52-Side | M_Pro7-Main | 12.99% |
|  |  |  |  | L_Arg52-Main | M_Ala8-Main | 10.43% |  |  |  | M_Ser-GalNAc-Side | H_Tyr101-Side | 11.43% |  |  |  | L_Arg52-Main | M_Ala8-Main | 10.51% |
| **16A** | Donor | Acceptor | Occupancy | Donor | Acceptor | Occupancy | Donor | Acceptor | Occupancy | Donor | Acceptor | Occupancy | Donor | Acceptor | Occupancy | Donor | Acceptor | Occupancy |
|  | M_Ala5-Main | H_Arg32-Main | 84.81% | M_Ala5-Main | H_Arg32-Main | 86.93% | M_Ala5-Main | H_Arg32-Main | 85.29% | H_Glu102-Main | M_Pro6-Main | 89.65% | M_Ala5-Main | H_Arg32-Main | 84.69% | M_Ala5-Main | H_Arg32-Main | 84.69% |
|  | H_Glu102-Main | M_Pro6-Main | 53.96% | H_Glu102-Main | M_Pro6-Main | 81.57% | H_Glu102-Main | M_Pro6-Main | 84.65% | M_Ala5-Main | H_Arg32-Main | 89.21% | H_Glu102-Main | M_Pro6-Main | 72.42% | H_Glu102-Main | M_Pro6-Main | 80.18% |
|  | L_Arg52-Side | M_Pro7-Main | 52.12% | L_Asn36-Side | M_Ala8-Main | 59.59% | L_Asn36-Side | M_Ala8-Main | 55.48% | L_Asn36-Side | M_Ala8-Main | 50.56% | H_Trp34-Side | M_Thr-GalNAc-Side | 33.65% | L_Asn36-Side | M_Ala8-Main | 48.68% |
|  | L_Arg52-Side | M_Gly10-Main | 46.92% | H_Arg32-Side | M_Ser3-Main | 36.73% | L_Arg52-Side | M_His9-Main | 30.30% | M_Thr-GalNAc-Side | H_Asp55-Side | 46.36% | L_Arg52-Side | M_His9-Main | 30.70% | H_Trp34-Side | M_Thr-GalNAc-Side | 39.09% |
|  | H_Arg32-Side | M_Ser3-Main | 36.69% | L_Arg52-Side | M_Gly10-Main | 23.10% | H_Arg32-Side | M_Ser3-Main | 25.98% | M_Ser-GalNAc-Side | H_Tyr101-Side | 28.62% | H_Arg32-Side | M_Ser3-Main | 23.22% | L_Arg52-Side | M_His9-Main | 33.09% |
|  | M_His9-Side | H_Glu51-Side | 12.11% | L_Arg52-Side | M_His9-Main | 21.98% | M_Ser-Gal(Core1)-Side | H_Asp29-Side | 17.55% | L_Arg52-Side | M_Pro7-Main | 24.14% | L_Arg52-Side | M_Pro7-Main | 20.74% | H_Arg32-Side | M_Ser3-Main | 19.70% |
|  |  |  |  | M_Ser-GalNAc-Side | H_Tyr101-Side | 15.83% | L_Arg52-Side | M_Pro7-Main | 15.63% | H_Trp34-Side | M_Thr-GalNAc-Side | 18.19% |  |  |  | M_Thr-GalNAc-Side | H_Asp55-Side | 17.63% |
|  |  |  |  | H_Trp34-Side | M_Ala5-Main | 14.27% | H_Arg32-Side | M_Gly2-Main | 11.87% | L_Arg52-Side | M_His9-Main | 17.79% |  |  |  | H_Arg32-Side | M_Gly2-Main | 14.75% |
|  |  |  |  | L_Arg52-Side | M_Pro7-Main | 10.83% | H_Arg32-Side | M_Pro1-Main | 10.07% | H_Arg32-Side | M_Ser3-Main | 16.95% |  |  |  | H_Arg32-Side | M_Pro1-Main | 10.91% |
|  |  |  |  |  |  |  |  |  |  | H_Arg32-Side | M_Pro1-Main | 15.31% |  |  |  | L_Arg52-Side | M_Pro7-Main | 10.67% |
|  |  |  |  |  |  |  |  |  |  | H_Arg32-Side | M_Gly2-Main | 11.39% |  |  |  |  |  |  |

* “H_” represents heavy chain of antibody, “L_” represents light chain of antibody, and “M_” represents MUC1-peptides. “Side” means the hydrogen bond formed by the atoms in side chain of residue, and “Main” means formed by the atoms in main chain.
